# Metamorphosis reverses the behavioural phenotype in *Rana arvalis* along a latitudinal gradient

**DOI:** 10.1101/2023.10.13.562178

**Authors:** Maria Cortazar-Chinarro, Alberto Corral-Lopez, Deike U. Lüdtke, Fredrik Tegner, Emilien Luquet, Anssi Laurila

**Affiliations:** Department of Ecology and Genetics, Uppsala University, Norbyvägen 18D, 75236 Uppsala, Sweden; ISOE—Institute for Social-Ecological Research, 60486 Frankfurt, Germany; Senckenberg Biodiversity and Climate Research Centre SBiK-F, 60325 Frankfurt/Main, Germany; Université Claude Bernard Lyon 1, LEHNA UMR 5023, CNRS, ENTPE, F-69622, Villeurbanne, France; Institut universitaire de France (IUF)

**Keywords:** amphibian, dual post-glacial recolonization route, POLS, pace-of-life

## Abstract

1. Understanding how demographic processes and environmental conditions shape behavioural variation across populations is pivotal in evolutionary ecology. However, the role that such processes play in the link between behaviour and life-history traits across populations remains largely unclear.
2. The moor frog (*Rana arvalis*) has colonized Sweden via two distinct routes from south via Denmark and from north via Finland. We collected *R. arvalis* eggs from multiple populations along a 1700 km latitudinal gradient across Northern Europe and raised tadpoles in a common garden experiment. We assessed developmental growth and proactivity levels in ca. 300 individuals at two key stages of anuran larval development: tadpoles (Gosner stage 32) and froglets (Gosner stage 46).
3. We found strong behavioural differences along the latitudinal gradient and between developmental stages. Tadpoles from northernmost populations were bolder (shorter times to leave a shelter) and showed higher activity levels in an open field test compared to those from southern populations.
4. However, these behavioural patterns reversed at the froglet stage, with individuals from northern populations showing reduced proactivity that those from southern populations. Further analyses indicated significant associations between developmental growth and boldness, with contrasting patterns across developmental stages and colonization routes.
5. In line with recent revisitations of the pace-of-life syndrome theory (POLS) for animal personality, our findings suggest a decoupling of correlations between behaviour and life history traits across ontogeny, likely reflecting adaptive responses to divergent ecological and demographic constraints along the latitudinal gradient.

## INTRODUCTION

Organisms display astonishing variation in behavioural and life history traits (Healy et al., 2019; Stearns, 1992; Wilson et al., 1994; Wolf, 2007). The Pace-Of-Life Syndrome (POLS) has been proposed to explain how the continuum of proactive-reactive behaviours is expected to coevolve with specific physiological and life history traits including metabolic rate, fecundity and growth (Biro and Stamps, 2008; Careau et al., 2008; Réale et al., 2010; Stamps, 2007). In POLS, trade-offs between life history, physiology and behaviour are hypothesized to lead to consistent inter-individual differences in behaviour across time and contexts—often referred to as personality (Bakker, 1986; Bouchard and Loehlin, 2001; Groothuis and Carere, 2005; Wilson et al., 1994). The POLS framework has given valuable insights on how animal personalities may constrain individual responses to environmental variation (e.g. Spiegel et al., 2015, Villegas-Rios et al., 2018) and has been successful in explaining variation especially at species level (e.g. Healy et al., 2019; van de Valle et al., 2023). However, mixed empirical evidence at within-species level suggests a need to integrate additional factors, such as species-specific demographic history and ecological pressures, to better understand the links between behaviour, physiology, and life history (e.g., Chang et al., 2024; Gopal et al., 2023; Montiglio et al., 2018).

Latitudinal gradients represent one of the most important biogeographic patterns on the Earth (Fischer, 1960). Populations at higher latitudes face harsher climatic conditions and shorter breeding seasons - environmental pressures that also influence the composition of predator and parasite communities and intensity of biotic interactions (Willig et al., 2003; Poulin and Leung, 2011; Schemske et al., 2009). Consequently, environmental conditions at high latitudes drive the evolution of distinct life-history strategies (Stearns 1992). Across large-scale latitudinal gradients, POLS predicts a different pace-of-life along gradients, with populations at higher latitudes showing more proactive, risk-prone behaviours (Cassidy et a.l, 2020; Foster et al., 2015; Gerlai and Csányi, 1990; Higgins et al., 2022; Mitchell et al., 1977). For example, in many anuran tadpoles, more stringent climatic gradient at high altitudes favors faster growth and development rates (Berven et al., 1983; Laugen et al., 2003, Orizaola et al., 2010, Luquet et al., 2019). Faster growth and development rates are linked with higher foraging activity which is likely allowed by lower predator density at high latitudes (Laurila et al., 2006, 2008). Similar relationships have been observed in other environmental contexts (e.g., between island and continental populations: Brodin et al., 2013; along altitudinal gradient: Luquet et al., 2015). However, other studies challenge this pattern indicating a complex relationship between latitude and personality traits. For example, populations of eastern mosquitofish, *Gambusia holbrooki,* show decreased proactivity at high-latitude populations with lower predation pressure (Culumber, 2022). To address these inconsistencies and gain a deeper understanding of how geographical variation influences behaviour and life-history strategies, further research is needed.

Most investigations of POLS focus on individual behaviours at a single life stage (Cabrera et al 2021). However, personality variation may emerge across developmental stages in populations exposed to diverse ecological conditions. This gap should be of particular importance in organisms showing complex life cycles such as amphibians. These species exhibit a dramatic metamorphosis, involving substantial changes in morphology, physiology and behaviour as aquatic larvae transition into terrestrial juveniles (Cohen, 1985; Wilbur, 1980). Consequently, amphibian larvae and juveniles vary greatly in many ecological aspects, for example by occupying distinct trophic niches (Kelleher et al., 2018; Rowe and Ludwig, 1991; Wilbur, 1980). Most anuran larvae are herbivores or omnivores, while post-metamorphic individuals are semi-aquatic carnivores, with dispersal and reproduction occurring at this stage (Wilbur, 1980). The shift of habitat at metamorphosis may also lead to substantial change in predation pressure (Alford, 1999). Under the POLS framework, selection over such differentiated ecological conditions should act to uncouple behavioural syndromes across ontogenetic stages in amphibians (Sih et al., 2004). Previous studies in species with complex life cycles have found both significant consistency (Begue et al., 2024; Brodin 2009; Wilson and Krause 2012) and nonsignificant/weaker relationships (Amat et al., 2018; Brodin et al., 2013; Moncaeu et al., 2017; Plaskonka et al., 2024) in proactivity levels over the metamorphic boundary. This discrepancy between theoretical works and empirical results may result from the lack of consideration of the relationship between personality and life history according to ecological conditions. Consequently, understanding the variation of behaviour in the proactive-reactive continuum across developmental stages in amphibian populations and how they associate to life history across multiple ecological conditions is crucial to resolve this apparent discrepancy.

North European populations of the moor frog (*Rana arvalis*) provide a suitable model for investigating evolutionary processes shaping behavioural phenotypes and life-history traits across diverse ecological conditions. This species has a wide latitudinal distribution with well-documented variation in life history traits along this geographical range (e.g., Laurila et al., 2006; Luquet et al., 2019; Räsänen et al., 2008). Following the last glaciation, the species colonized the Scandinavian Peninsula through two distinct routes: one from the north via present-day Finland and another from the south, with a contact zone established in northern central Sweden (Cortazar-Chinarro *et al*. 2017; Knopp and Merilä, 2009,). These post-glacial colonization events and subsequent demographic processes have significantly influenced the divergence between populations. Luquet et al. (2019) demonstrated strong divergence between southern and northern lineages, with strong selection pressures driving higher growth, development rates and larger body size in northern populations. Additionally, genetic analyses revealed clear differentiation between two major genetic clusters corresponding to northern and southern populations (Cortazar-Chinarro *et al.,* 2017; Meyer-Lucht *et al*., 2019, Rödin-Mörch *et al*., 2019).

In this study, we evaluate behavioural variation across populations, regions and genetic clusters of *R. arvalis* along a latitudinal gradient, incorporating information from two developmental stages. We conducted a common garden experiment using populations from five different regions in northern Europe and performed repeated open field tests on individuals before and after metamorphosis to quantify behavioural traits on the proactive-reactive continuum. This data was combined with life-history measurements to provide insight into the relationship between behaviour, development and life-history across large-scale geographical variation.

In *R. arvalis*, shorter breeding seasons and harsher climatic conditions at higher latitudes are associated with faster development (Luquet *et al*., 2019; Meyer-Lucht *et al*., 2019). Based on the POLS framework, we hypothesized that i) proactive behaviours would be positively selected in high latitudes at early developmental stages, resulting in a gradient of increasing proactivity-associated traits with increasing latitude; ii) behavioural response on the proactive-reactive continuum would be decoupled across ontogeny due to distinct ecological challenges and selective pressures faced during larval and post-metamorphic stages; and iii) proactive behaviours would positively coevolve with faster developmental growth throughout our geographical range.

## METHODS

### Sample Collection and latitudinal gradient

*R. arvalis* has a broad Eurasian distribution, from the North Sea coast to western Siberia (Sillero et al., 2014). We collected 50-100 eggs from each of 10 freshly laid clutches (hereafter families) at each of three breeding populations within five regions along a 1700 km latitudinal gradient from northern Germany (Hanover) to northern Sweden (Luleå), covering a major part of the latitudinal distribution of the species (more details about field sampling in Luquet et al., 2019). However, eggs from only one population were collected in Hanover. The timing of breeding season along the latitudinal gradient varies from mid-March (Hanover) to late May (Luleå). The distance between populations within the same region varied from 8 to 50 km (average 20 km). The eggs were transported to the laboratory in Uppsala and kept in a climate room at 16°C and 16L:8D photoperiod until they hatched and reached Gosner stage 25 (independent feeding; Gosner 1960).

### Experimental design

#### Common garden experiment

Tadpoles were reared in a common garden experiment as described in Luquet *et al*. (2019), however, the individuals included in this study were all reared at the 16 °C temperature treatment. At Gosner stage 25, 3 tadpoles from each of the 10 families per population were haphazardly selected. Tadpoles were raised individually in 0.75 L opaque plastic vials until metamorphosis. They were fed with chopped spinach *ad libitum* and the water was changed every third day. The experiment was conducted in walk-in climate-controlled rooms with 16 °C average water temperature, corresponding to a temperature commonly encountered in the natural environment. The photoperiod was 16L:8D, corresponding to the situation in the middle of the gradient (Uppsala) in May. When tadpoles reached Gosner stage 32 (Gosner, 1960), we performed the first set of behavioural assays at tadpole stage (see below), weighed them to the nearest 0.1 mg and returned them in their vials until they reached metamorphosis (emergence of the first forelimb; stage 42, Gosner, 1960). At metamorphosis, the water was removed and the vials were closed with a lid. On the day when the metamorphs reached the froglet stage (i.e. Gosner stage 46, tail entirely resorbed), we performed a second set of behavioural assays and weighed them to the nearest 0.1 mg.

#### Recording of behaviour

We recorded the behaviour of tadpoles and froglets (n tadpole / froglet: Hanover = 31 / 20, Skåne = 89 / 65, Uppsala = 90 / 73, Umeå = 89 / 65, Luleå = 90 / 61, total = 389 / 284) individually using an open field test in circular arenas (diameter = 90 cm) with a circular plastic shelter (diameter = 5 cm) in the center in a room at 16 °C. In the tadpole tests the arenas were filled with 10 cm of water while in froglet tests the arena floor was moist. Individuals were put in the shelter and left for 10 min to acclimate. The recording started when we opened the shelter. Videos were recorded by a digital videocamera (Sony, HDR-HC1E Handy Camera) placed 1.70 m above the experimental arena. Using the recordings, we scored emergence time from the shelter in an open field test, a widely used proxy for *boldness* (Mazué *et al*, 2015), by scoring the time point when the body of the tadpole or froglet had completely emerged from the shelter provided. In cases where the individual did not leave the shelter, the trial was stopped after 20 min for tadpoles (n = 6) and 30 min for froglets (n = 126).

For individuals that left the shelter we analyzed 5 minutes of recording to measure proactivity from two additional traits: i) *activity,* obtained as the mean speed during the recording and ii) *exploration* obtained as the average distance to the shelter during the recorded time. Videos were analyzed using the computer vision software Ctrax v0.5.19 (Branson *et al*, 2009) and the quality of the tracking was checked for every individual. When necessary, the tracking was corrected manually using the ‘fixerrors’ function. We then collapsed the frame-by-frame positional data obtained and calculated the behavioural proxies using the tracker package (https://github.com/Ax3man/trackr) in R v. 4.1.3 (R Core team 2024).

### Statistical analysis

#### Consistency of individual behaviour (personality)

To assess personality, i.e. behavioural consistency at the individual level, we recorded the behaviour of a representative subset of individuals for a second time 24-48 hours later (n tadpole / froglet: Hanover = 2 / 2, Skåne = 7 / 6, Uppsala = 6 / 8, Umeå = 6 / 8, Luleå = 9 / 9, total = 30 / 33). To avoid potential biases introduced by habituation to the experimental setup, individuals that were tested twice as tadpoles were not used for repeated measurements during froglet stage. As we could not obtain repeated measurements of activity or exploration for froglets that did not leave the shelter during the first or second trial (n = 17), we used data on time to leave the shelter to assess the individual repeatability of boldness. Repeatability, the relative partitioning of variance into within-group and between-group sources of variance (Dingemanse and Dochtermann, 2013; Nakagawa and Schielzeth, 2010) was estimated within a linear-mixed model (LMM) framework. We assessed the variance explained by individual identity in boldness (log-transformed emergence time) in independent gaussian models for tadpoles and froglets that included region as fixed effects, and population and individual identity as random factors using the ‘rptR’ package in Rv4.4.1 (Stoffel et al. 2017, R Core team 2024).

#### Behavioural traits across the latitudinal gradient

We assessed potential differences in behaviour across genetic clusters or region of origin and between developmental stages implementing LMMs in ‘lme4’ package in Rv4.4.1 (Bates et al. 2015; R Core Team 2024). Each behaviour measured was assessed in an independent model. For models to assess differences by region, developmental stage, region and their interaction were included as a fixed effect, and population of origin and individual identity as random intercept effects. For models to assess differences by genetic cluster, developmental stage, cluster and their interaction were included as a fixed effect, and population of origin, region and individual identity as random intercept effects. Individual mass at the time of recording was added as a covariate in all models. Model fit and significance were assessed through conditional F-tests with Kenward-Roger approximation for the degrees of freedom using pbkrtest and lmerTest R packages (Halekoh and Højsgaard 2014, Kutznesova et al. 2017). To assess differences between regions, we obtained post-hoc contrasts of the best fitted model using the ‘emmeans’ package and including false discovery rate correction for multiple tests (Lenth 2018). Significance of random effects was tested using likelihood ratio tests comparing models with or without the random effects in the full fixed effect structure.

A high number of froglets did not leave the shelter and were assigned to an emergence value of the decided cutoff point (1800 sec; n=126). Thus, time to emerge from shelter (boldness) data was analyzed using a mixed effect survival test performed with the ‘coxme’ package (Therneau 2015). The model structure was analogous to the LMMs above. Significance of fixed and random effects was tested using likelihood ratio tests comparing models with or without the tested effect in the full fixed effect structure.

We used repeated measurements to same individuals taken during tadpole and froglet stage to estimate the repeatability of behaviours over the metamorphic boundary using analogous LMMs than described above and modelling our data using the ‘rptR’ package in Rv4.4.1 (Stoffel et al. 2017, R Core team 2024).

#### Link between behaviour and life-history across genetic clusters

We combined data on time to reach metamorphosis (Gosner stage 42, in days) and mass at the same stage to calculate developmental growth (mass/time at Gosner stage 42) for the individuals evaluated for their behaviour. This life-history trait quantified for the individuals used in this behavioural study are a subset of data included in a study on the quantitative differentiation of life-history traits along the latitudinal gradient (Luquet *et al*, 2019; Meyer-Lucht *et al*, 2019). Hence, analyses on differences in life-history traits across the gradient are omitted in this study and we refer to these studies for more details of their differences across the populations sampled.

To assess the relationship between behaviour and life-history across genetic clusters and developmental stages, we structured LMMs testing the influence of behavioural traits on growth rate. Behavioural traits (i.e. boldness / activity / exploration, considered independently), genetic cluster or region and developmental stage were fixed effects and population of origin and individual identity were included as random intercept effects. Models fit and significance were assessed as previously through conditional F-tests with Kenward-Roger approximation for the degrees of freedom using pbkrtest and lmerTest R packages (Halekoh and Højsgaard 2014, Kutznesova et al. 2017). We obtained post-hoc contrasts of fitted models using the ‘emmeans’ package and including false discovery rate correction for multiple tests (Lenth 2018). Significance of random effects was tested using likelihood ratio tests comparing models with or without the random effects in the full fixed effect structure.

## RESULTS

### Repeatability of emergence time (boldness)

The time to emerge from the shelter was repeatable (controlling for the effects of region) at froglet stage, but not at tadpole stage (Tadpole: *R* = 0.000 ± 0.060 *SE*, *P* = 1, Froglet: *R* = 0.362 ± 0.189 *SE*, *P* = 0.023; Fig 1).

**Figure 1.**
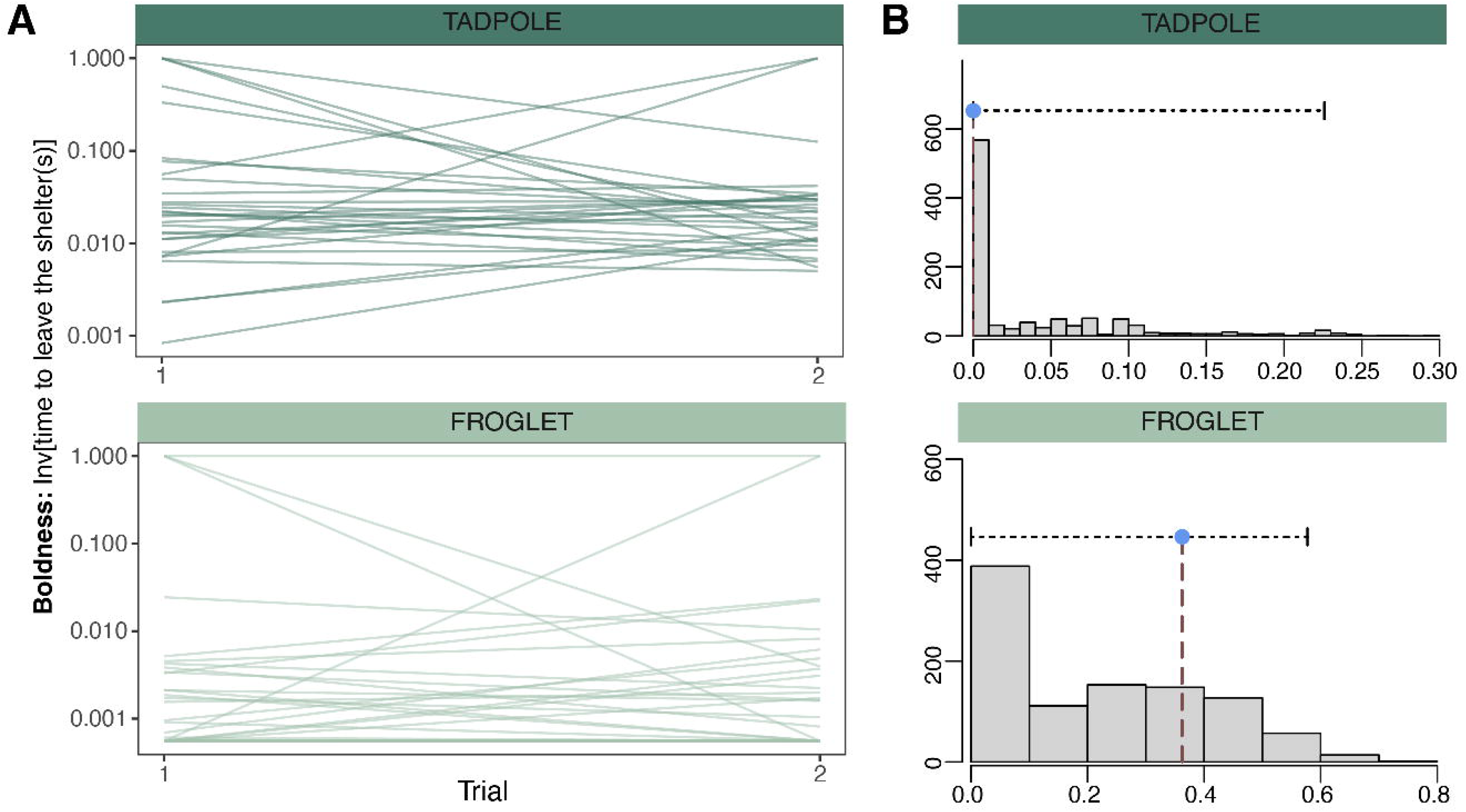
Consistency of individual behaviour. (A) Individual differences in measurements of time to emerge from the shelter taken during two independent trials performed to moor frog tadpoles (n = 30, dark green) and froglets (n= 33, light green) raised in a common garden experiment sampled across multiple regions of a 1700km latitudinal gradient in northern Europe. (B) Distribution of adjusted repeatability estimates of an individual time to emerge from the shelter in bootstrapping tests using a linear mixed model framework after controlling for effects introduced by region (see Stoffel et al. 2017). Blue dot indicates mean adjusted repeatability with confidence intervals for time to emerge in tadpoles (top) and froglets (bottom).

### Behaviour across genetic clusters, regions and developmental stages

#### Boldness

The time to emerge from the shelter was significantly influenced by the interaction between developmental stage and genetic cluster (Table 2). Specifically, boldness did not differ between tadpoles from the South and North genetic clusters, but froglets from South populations were bolder than froglets from North populations (Tadpole _North vs South_ ratio = 1.26 ± 0.21 *SE, z* = 1.40, *P* = 0.16, Froglet _North vs South_ ratio = 0.50 ± 0.08 *SE, z* = -4.47, *P* < 0.001; Fig 2A). Similarly, statistical models incorporating the region of origin indicated a significant interaction effect between developmental stage and region on time to emerge from the shelter (Table 3). Tadpoles from the northernmost region (Luleå) were significantly bolder than those from all the other regions sampled, including those from Umeå within the same genetic cluster (ratio [SE]: Hanover-Luleå: 0.38 [0.09] - *adjP < 0.001*; Skåne-Luleå: 0.65 [0.12] - *adjP = 0.040*; Uppsala-Luleå: 0.55 [0.09] - *adjP = 0.001*; Umeå-Luleå: 0.51 [0.08] - *adjP < 0.001;* Figure. 1D; Table S1). Tadpoles from the southernmost region (Hanover) tended to be shyer (longer time to emerge from the shelter) than those from the other regions although the difference was only significant in comparison to Skåne (ratio [SE]: 0.58 [0.13] - *adjP = 0.040;* Figure. 1D; Table S1). In froglets, individuals from the two northernmost regions were significantly shyer than individuals from the rest of regions studied in the gradient (Figure 1D, see Table S1). The random population effect was never significant for boldness (Tables 2 and 3 for models incorporating genetic clusters or regions, respectively). Boldness was not repeatable across ontogeny (i.e. between tadpole and froglet stages; *R* = 0.079 ± 0.056 *SE*, *P* = 0.10; Figure S1).

**Table 1.** Consistency of individual behaviour. Results for repeatability analyses of measurements of time to emerge from a shelter taken twice to moor frogs during tadpole stage ( n = 30), and froglet stage ( n= 33) raised in a common garden experiment following collection from multiple regions across a 1700km latitudinal gradient in northern Europe. Using a linear mixed model framework, we estimated adjusted repeatability of boldness for individual and population after controlling for effects introduced by region in the model (_adj_R; see Stoffel et al. 2017), as well as the quantification of uncertainty for the variance explained by region of origin (R^2^ coefficient, see Stoffel et al. 2017).

| Developmental stage | Factor | $_{adj}R / R^2$ | *SE* | 95% CI | P |
| --- | --- | --- | --- | --- | --- |
| Tadpole | Individual | 0.00 | 0.06 | [0, 0.226] | 1.00 |
|  | Population | 0.00 | 0.06 | [0, 0.131] | 1.00 |
|  | Fixed (region) | 0.23 | 0.16 | [0.088, 0.683] | - |
| Froglet | Individual | 0.36 | 0.19 | [0, 0.577] | **0.023\*** |
|  | Population | 0.00 | 0.18 | [0, 0.535] | 1.00 |
|  | Fixed (region) | 0.24 | 0.18 | [0.079, 0.769] | - |

**Table 2.** Behavior across genetic clusters and developmental stages. Results for statistical analyses evaluating behavioral traits across tadpole and froglet stages in moor frogs originated from five different regions across a 1700 latitudinal gradient and raised in a common garden experiment (South genetic cluster: Hanover (Germany), Skåne (Sweden), Uppsala), North genetic cluster: Umeå (Sweden), Luleå (Sweden)). Boldness data was analyzed using a mixed effect survival model, while activity and exploration data was analyzed using linear mixed models. Full models included genetic cluster, developmental stage and interactions as fixed effects, while population of origin, region and individual identity as random intercept effects. Individual mass at the time of recording was added as a covariate.

| **Boldness:** Time to leave the shelter |  |  |  |  | **Activity:** mean speed |  |  |  | **Exploration:** mean distance to shelter |  |  |  |
| --- | --- | --- | --- | --- | --- | --- | --- | --- | --- | --- | --- | --- |
| *Predictors* | $\Delta \log\text{-lik}$ | $\chi^2$ | *p* | *df* | *SumSq*<br>(mean sq) | *F-value* | *p* | *df (Num<sub>dem</sub>)* | *SumSq*<br>(mean sq) | *F-value* | *p* | *df (Num<sub>dem</sub>)* |
| cluster | 16.0 *** | 32.04 | **<0.001** | 1 | 0.04<br>(0.04) | 1.33 | 0.32 | 1<sub>3.23</sub> | 785.1<br>(785.1) | 6.02 | 0.064 | 1<sub>4.42</sub> |
| stage | -31.5 *** | 137.03 | **<0.001** | 1 | 1.96 ***<br>(1.96) | 61.07 | **<0.001** | 1<sub>466.5</sub> | 716.5 *<br>(716.5) | 5.50 | **0.019** | 1<sub>412.8</sub> |
| mass at recording | 2.6 * | 5.22 | **0.022** | 1 | 0.14*<br>(0.13) | 4.23 | **0.040** | 1<sub>504.3</sub> | 157.0<br>(157.0) | 1.21 | 0.27 | 1<sub>409.7</sub> |
| cluster × stage | 14.5 *** | 28.98 | **<0.001** | 1 | 1.10***<br>(1.10) | 34.26 | **<0.001** | 1<sub>310.6</sub> | 1164.6**<br>(1164.6) | 8.94 | **0.003** | 1<sub>339.9</sub> |
| **Random Effects** | $\Delta \log\text{-lik}$ | $\chi^2$ | *p* | *df* | $\Delta \log\text{-lik}$ | $\chi^2$ | *p* | *df* | $\Delta \log\text{-lik}$ | $\chi^2$ | *p* | *df* |
| id | 0.0 | 0.00 | 0.97 | 1 | -1.4 | 2.77 | 0.096 | 1 | 0.0 | 1.81 | 0.18 | 1 |
| population | 0.8 | 1.66 | 0.19 | 1 | 3.0 * | 6.07 | **0.013** | 1 | 0.0 | 0.0 | 1 | 1 |
| region | 0.0 | 0.05 | 0.81 | 1 | 0.2 | 0.55 | 0.45 | 1 | 0.0 | 0.13 | 0.71 | 1 |
| $\sigma^2$ | | | | | 0.03 | | | | 130.28 | | | |
| $\tau_{00}$ | 0.00 id | | | | 0.01 id | | | | 14.62 id | | | |
|  | 0.01 population |  |  |  | 0.00 population |  |  |  | 0.00 population |  |  |  |

|  | 0.00 region | 0.00 region | 2.09 region |
| --- | --- | --- | --- |
| Observations | 655 | 512 | 512 |
| Marginal R<sup>2</sup> | 0.71 | 0.365 | 0.058 |

*\*p<0.05 \*\*p<0.01 \*\*\*p<0.001*

**Figure 2.**
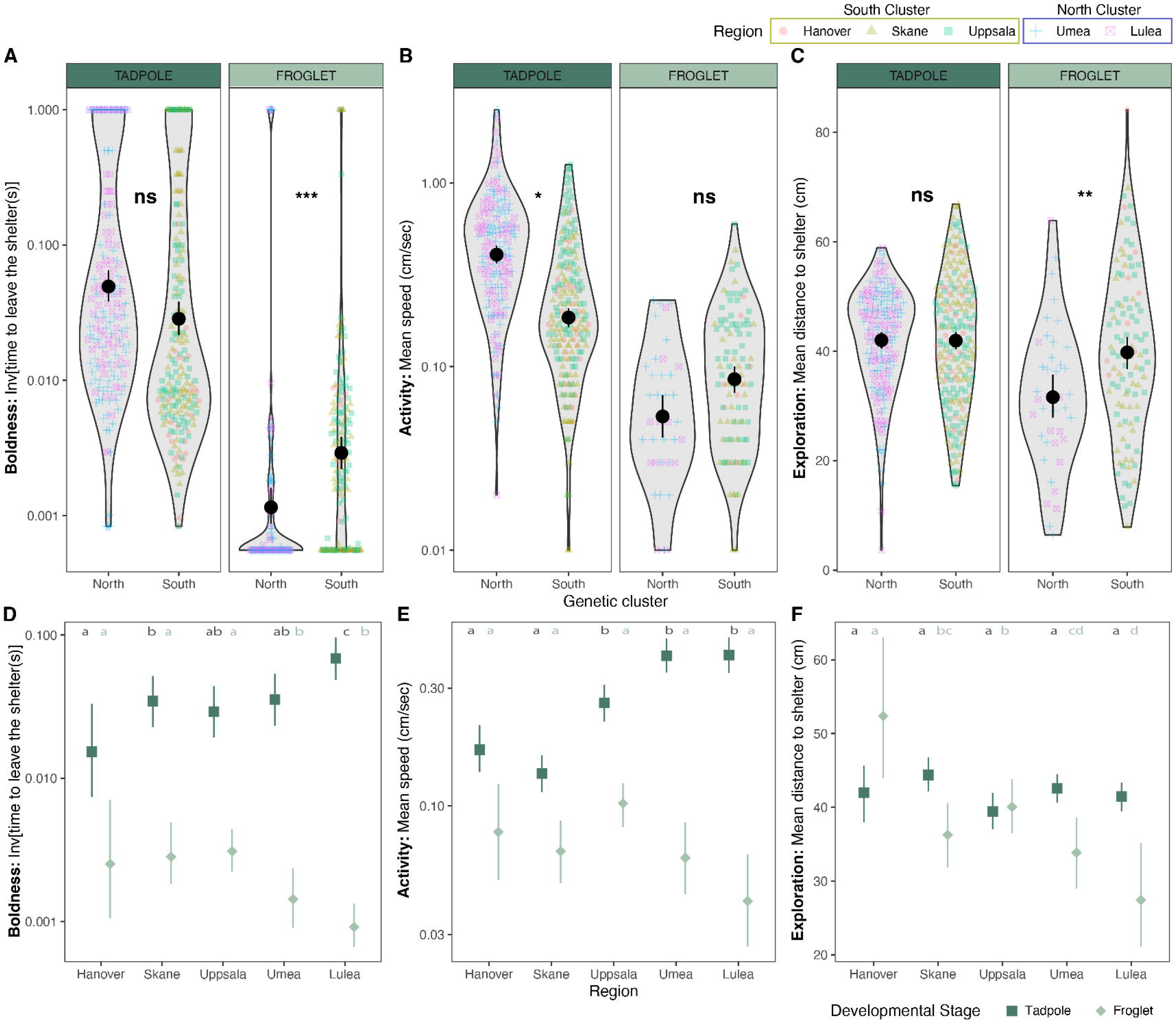
Behaviour of moor frogs across the latitudinal gradient. Boldness (A), activity (B) and exploration (C) measured in moor frog tadpoles (dark green) and froglets (light green) across a 1700 latitudinal gradient and raised in a common garden experiment (South genetic cluster: Hanover (Germany), red dot; Skåne (Sweden), yellow triangle; Uppsala, green square); North genetic cluster: Umeå (Sweden), blue cross; Luleå (Sweden), pink crossed square). Text indicates p-values of significance tests comparing difference in behaviour between genetic clusters (see Table 2; ns p>0.05, * p<0.05 ** p<0.01 *** p<0.001). Mean value and standard error of boldness (D), activity (E) and exploration (F) of tadpoles (dark green squares) and froglets (light green diamonds) for the five regions sampled. Average values for regions not sharing any letter are significantly different (p < 0.05) in post-hoc contrasts of independent models assessing differences in behaviour across regions for tadpoles (dark green letters) and froglets (light green letters; see Table 3-4).

**Table 3.** Behavior across regions and developmental stages. Results for statistical analyses evaluating behavioral traits across tadpole and froglet stages in moor frogs originated from five different regions across a 1700 latitudinal gradient and raised in a common garden experiment (Hanover (Germany), Skåne (Sweden), Uppsala), Umeå (Sweden), Luleå (Sweden)). Boldness data was analyzed using a mixed effect survival model, while activity and exploration data was analyzed using linear mixed models. Full models included developmental stage, region and its interaction as fixed effects, while population of origin, and individual identity as random intercept effects. Individual mass at the time of recording was added as a covariate.

| *Predictors* | **Boldness:** Time to leave the shelter |  |  |  | **Activity:** mean speed |  |  |  | **Exploration:** mean distance to shelter |  |  |  |
| --- | --- | --- | --- | --- | --- | --- | --- | --- | --- | --- | --- | --- |
| | $\Delta \log\text{-lik}$ | $\chi^2$ | *p* | *df* | *SumSq*<br>(mean sq) | *F-value* | *p* | *df* (*Num<sub>dem</sub>*) | *SumSq*<br>(mean sq) | *F-value* | *p* | *df*<br>(*Num<sub>dem</sub>*) |
| region | 29.2 * | 58.58 | **<0.001** | 1 | 0.23<br>(0.06) | 1.80 | 0.20 | 4<sub>11.1</sub> | 2606.1 **<br>(651.5) | 5.23 | **0.007** | 4<sub>16.5</sub> |
| stage | 78.9 *** | 156.29 | **<0.001** | 1 | 1.53 ***<br>(1.53) | 48.05 | **<0.001** | 1<sub>453.4</sub> | 204.2<br>(204.2) | 1.65 | 0.20 | 1<sub>455.2</sub> |
| mass at recording | 2.3 * | 4.73 | **0.030** | 1 | 0.12 *<br>(0.12) | 3.91 | **0.048** | 1<sub>500.4</sub> | 163.7<br>(163.7) | 1.32 | 0.25 | 1<sub>450.1</sub> |
| region × stage | 24.3 *** | 48.66 | **<0.001** | 1 | 1.24 ***<br>(0.31) | 9.71 | **<0.001** | 4<sub>288.0</sub> | 3511.7 ***<br>(877.7) | 7.09 | **<0.001** | 4<sub>309.7</sub> |
| **Random Effects** | $\Delta \log\text{-lik}$ | $\chi^2$ | *p* | *df* | $\Delta \log\text{-lik}$ | $\chi^2$ | *p* | *df* | $\Delta \log\text{-lik}$ | $\chi^2$ | *p* | *df* |
| id | 0.0 | 0.00 | 0.95 | 1 | 1.6 | 3.25 | 0.071 | 1 | -0.8 | 2.40 | 0.12 | 1 |
| population | 0.0 | 0.08 | 0.77 | 1 | 1.4 | 2.74 | 0.098 | 1 | 0.0 | 0.00 | 1 | 1 |
| $\sigma^2$ | | | | | 0.03 | | | | 123.83 | | | |
| $\tau_{00}$ | 0.00<sub>id</sub> | | | | 0.01<sub>id</sub> | | | | 16.85<sub>id</sub> | | | |
|  | 0.00<sub>population</sub> |  |  |  | 0.00<sub>population</sub> |  |  |  | 0.00<sub>population</sub> |  |  |  |
| Observations | 655 |  |  |  | 512 |  |  |  | 512 |  |  |  |
| Marginal R<sup>2</sup> | 0.725 |  |  |  | 0.389 |  |  |  | 0.104 |  |  |  |

\* $p < 0.05$ \*\* $p < 0.01$ \*\*\* $p < 0.001$

#### Activity

Activity was significantly influenced by the interaction between developmental stage and genetic cluster (Table 2) or region (Table 3). At the genetic cluster level, tadpoles from South populations were less active than those from North populations, but there was no difference at froglet developmental stage (Activity: Tadpole _North vs South_ estimate = 0.18 ± 0.05 *SE, t* = 3.41, *P* = 0.036, Froglet _North vs South_: estimate = -0.06 ± 0.06 *SE, t* = -0.98, *P* = 0.36; Fig 2B). At the regional level, there were differences in activity of tadpoles among the regions while in contrast all froglets have similar activity. Specifically for the tadpole stage, the activity increased with latitude: tadpoles from the two southernmost regions (Hanover, Skåne) had a lower activity than tadpoles from intermediate latitude (Uppsala region) and northernmost regions (Umeå, Luleå); the activity of tadpoles from intermediate region (Uppsala) was marginally lower than those from northernmost regions (Fig 2E, see Table S1). The random population effect in activity was significant in genetic cluster model (Table 2) and only marginal in region model (Table 3). Repeated measurements across developmental stages indicated significant, repeatability of activity across ontogeny (*R* = 0.158 ± 0.078 *SE*, *P* = 0.034; Figure S1).

#### Exploration

Exploration was significantly influenced by the interaction between developmental stage and genetic cluster (Table 2) or region (Table 3). Exploration of tadpoles from the South and North genetic clusters was similar while froglets from the North genetic cluster showed reduced exploration than those from the Southern cluster (Exploration: Tadpole _North vs South_ estimate = -1.07 ± 2.09 *SE, t* = -0.51, *P* = 0.63, Froglet _North vs South_: estimate = -9.03 ± 2.76 *SE, t* = -3.27, *P* = 0.005; Fig 2C). Specifically, for the froglet stage, the exploration of the southernmost region (Hanover) was higher than all other regions along the gradient (Figure. 2F, Table S14). Froglets from the two northernmost regions (Luleå and Umeå) had significantly lower exploration than froglets from southern regions (except between Skåne and Umeå; Fig 2F, Table S1). The random population effect was never significant (Tables 2 and 3 for models incorporating genetic clusters or regions, respectively). Repeated measurements across developmental stages indicated no repeatability of exploration (*R* = 0.099 ± 0.078 *SE*, *P* = 0.13; Figure S1).

### Influence of behavioural traits on growth rate

#### Growth/boldness

Growth rate was affected by the three-way interaction between boldness, genetic cluster and developmental stage (Fig 3A, Table 4). In the South genetic cluster, tadpoles with fastest times to emerge from the shelter (bolder) showed faster growth rates, but this relationship was not found in tadpoles originating from the North cluster (Figure 3A, Tadpole _North vs South_ estimate = 0.74 ± 0.28 *SE*, *P* = 0.008,Table S2). This pattern was reversed during froglet stage, as bolder individuals from the North cluster showed faster growth rates as tadpoles, while no association for froglets was found in the South cluster (Figure 3A, Table S2). Similarly, statistical models incorporating the region of origin indicated a significant three-way interaction between boldness, region and developmental stage (Fig S2A, Table 5). This effect was driven by the positive association between growth rate and boldness in Uppsala (estimate _Tadpole_ = 0.79 ± 0.26 *SE*, *P* = 0.027) and Skåne (estimate _Tadpole_ = 0.89 ± 0.35 *SE*, *P* = 0.059; see Table S3). Additionally, it was driven by i) differential associations of growth rate/boldness associations of these two regions in contrasts with all other regions (see Table S4; Figure S2A); and ii) in contrast with this relationship in froglets in Uppsala (estimate _Froglet_ = -0.01 ± 0.07 *SE*, Tadpole vs Froglet: t = 2.91, *P* = 0.004; Table S5; FigureS2A) and Skåne (estimate _Froglet_ = 0.11 ± 0.07 *SE*, Tadpole vs Froglet: t = 2.16, *P* = 0.031; Table S5; Figure S2A). The random population effect in activity was significant in models incorporating genetic cluster or region (Tables 4 and 5 respectively).

**Figure 3.**
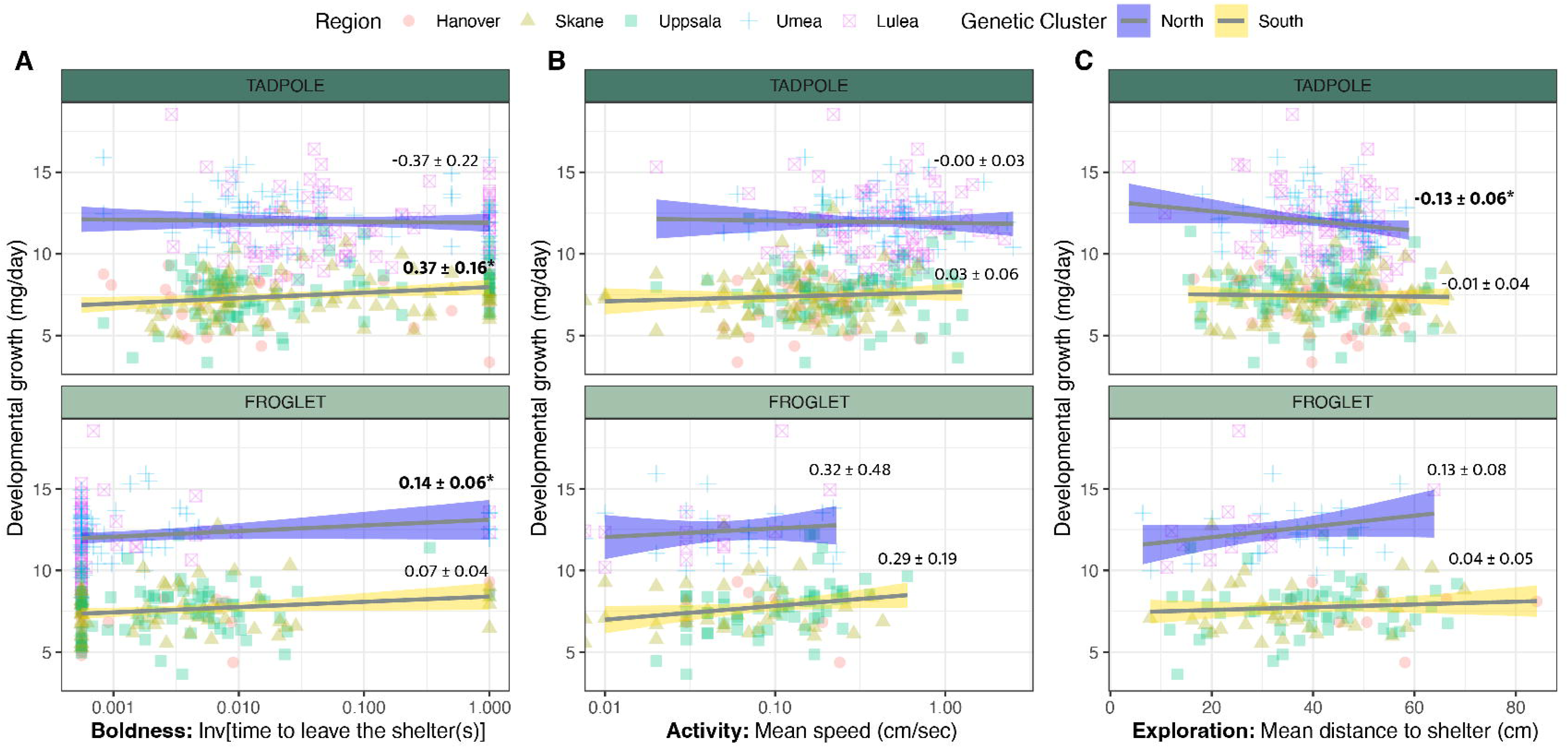
Association between behaviour and life history in moor frogs across the latitudinal gradient. Relationships between (A) boldness, (B) activity and (C) exploration and developmental growth (mass/time at Gosner stage 42) in moor frog tadpoles (top row) and froglets (bottom row) sampled across a 1700 latitudinal gradient and raised in a common garden experiment (South genetic cluster: Hanover (Germany), red dot; Skåne (Sweden), yellow triangle; Uppsala, green square); North genetic cluster: Umeå (Sweden), blue cross; Luleå (Sweden), pink crossed square). Trend lines and confidence interval are displayed for the relationship between each behaviour and developmental growth for values at the North (blue) and South (yellow) genetic clusters. Text indicates estimated marginal means and standard error for each trend in post-hoc contrasts of statistical models (p < 0.005 in bold; see Tables 5-6).

**Table 4.**
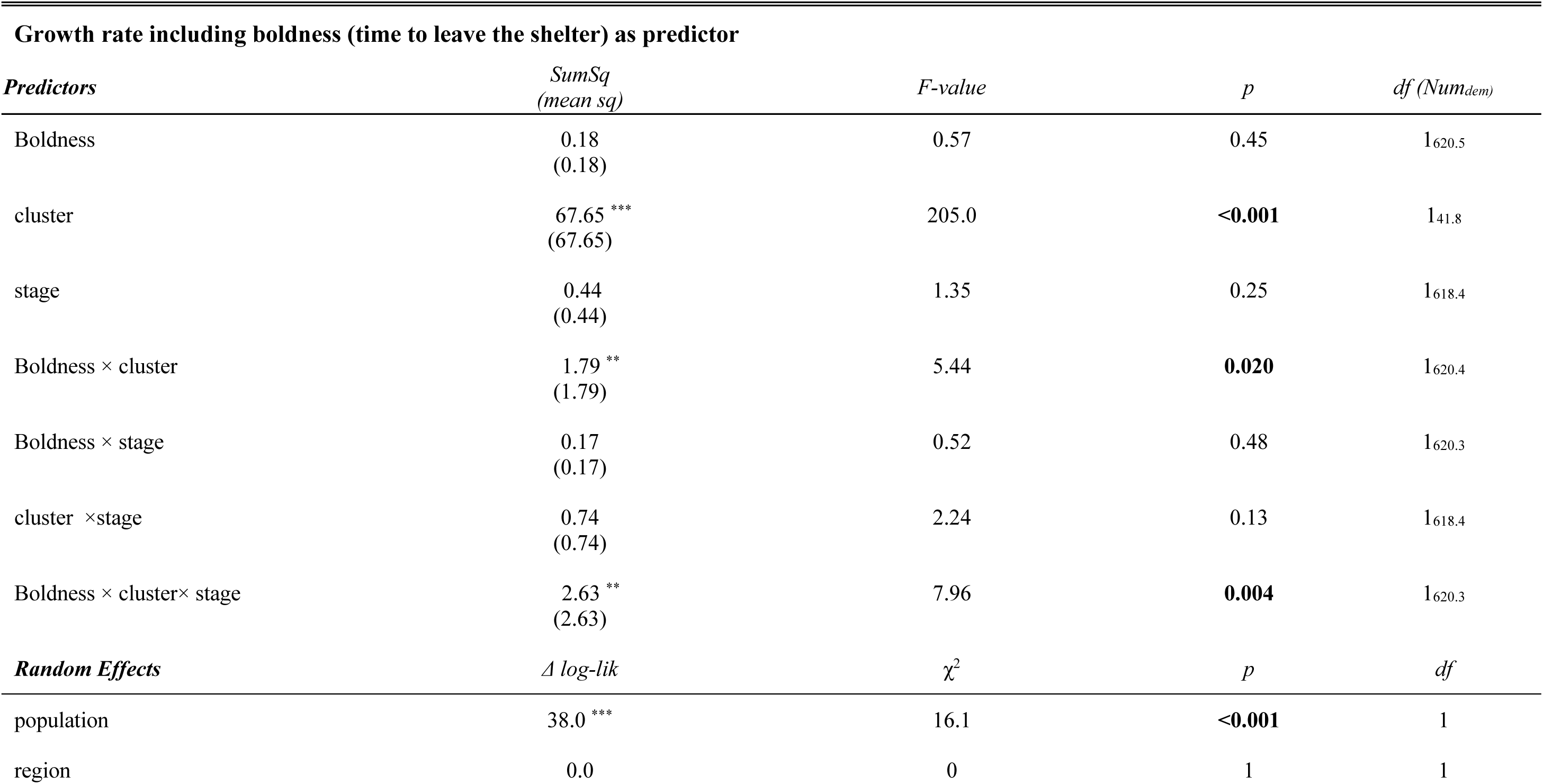
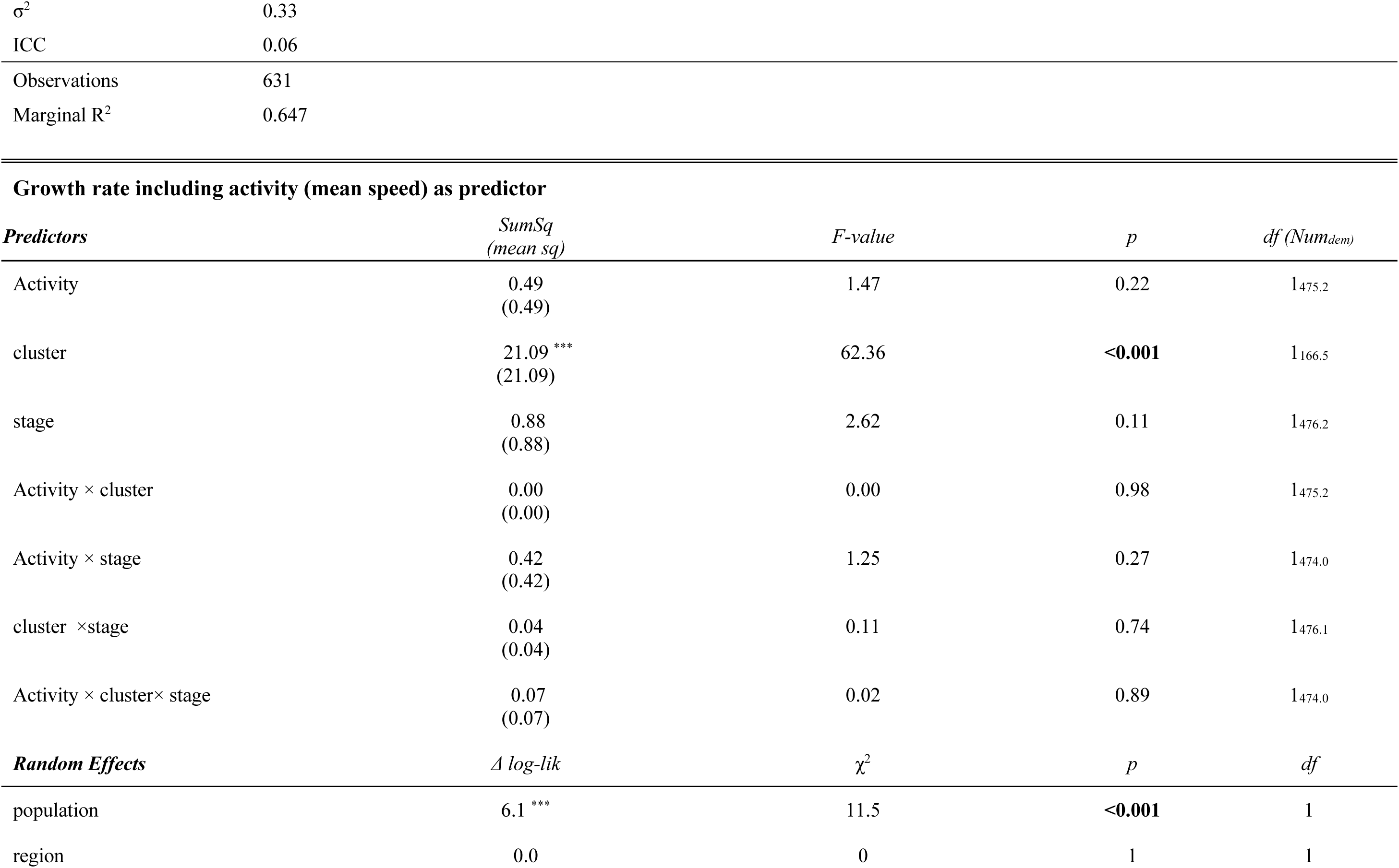
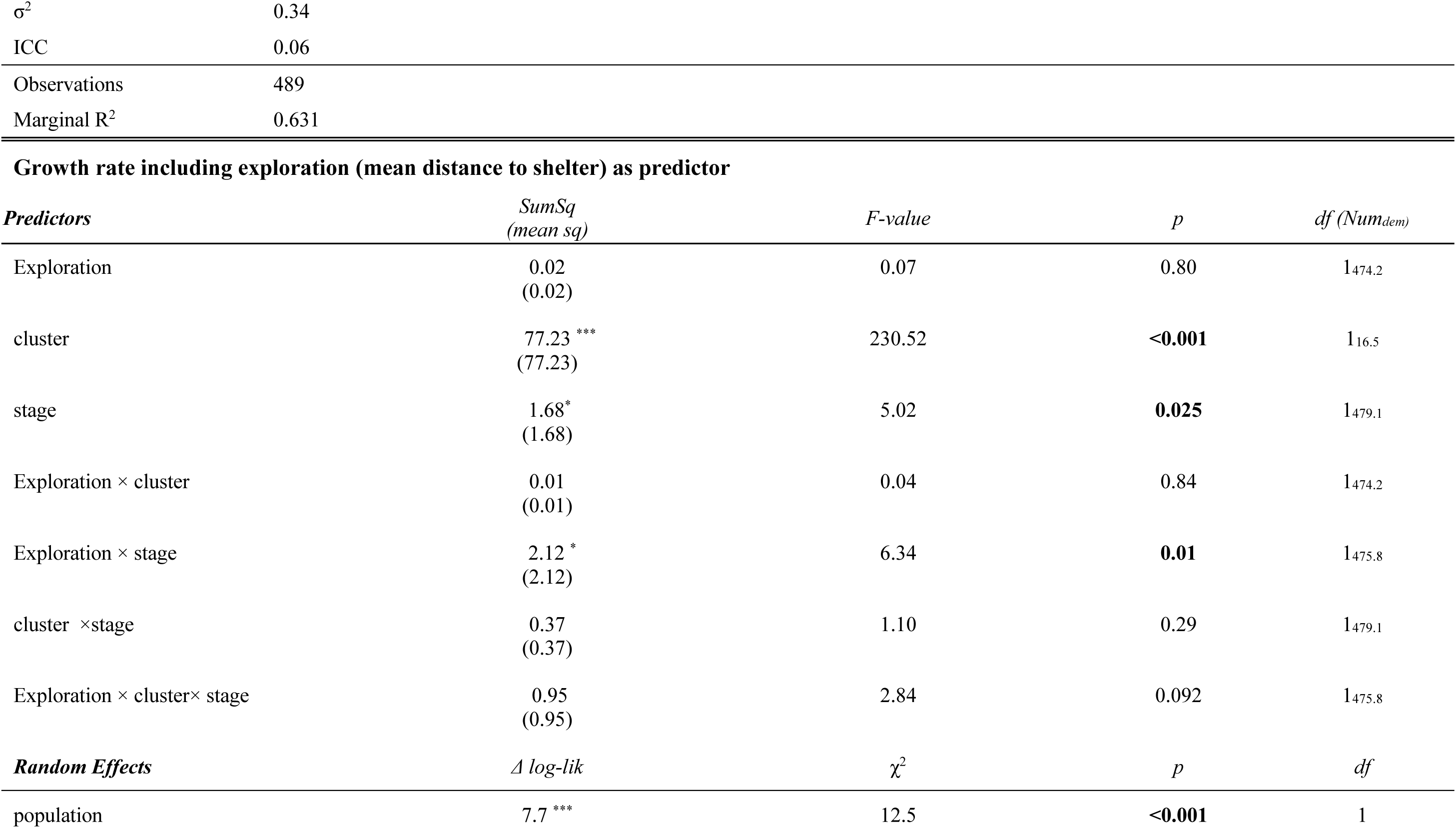
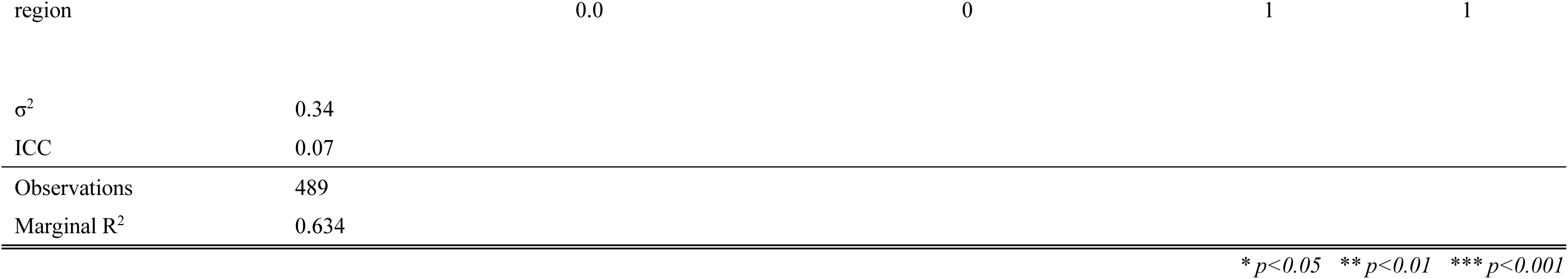
Influence of behavior on life history (growth rate) across genetic clusters and developmental stages. Results for statistical analyses evaluating growth rate in moor frog tadpoles and froglets originating from five different regions across a 1700 latitudinal gradient and raised in a common garden experiment (South genetic cluster: Hanover (Germany), Skåne (Sweden), Uppsala), North genetic cluster: Umeå (Sweden), Luleå (Sweden)). Growth rate linear mixed models included the behavior of interest, genetic cluster, developmental stage and their interaction as predictors, and population of origin and and region as random intercept effects.

| **Growth rate including boldness (time to leave the shelter) as predictor** |  |  |  |  |
| --- | --- | --- | --- | --- |
| *Predictors* | *SumSq*<br>*(mean sq)* | *F-value* | *p* | *df (Num<sub>dem</sub>)* |
| Boldness | 0.18<br>(0.18) | 0.57 | 0.45 | 1<sub>620.5</sub> |
| cluster | 67.65 ***<br>(67.65) | 205.0 | **<0.001** | 1<sub>41.8</sub> |
| stage | 0.44<br>(0.44) | 1.35 | 0.25 | 1<sub>618.4</sub> |
| Boldness × cluster | 1.79 **<br>(1.79) | 5.44 | **0.020** | 1<sub>620.4</sub> |
| Boldness × stage | 0.17<br>(0.17) | 0.52 | 0.48 | 1<sub>620.3</sub> |
| cluster × stage | 0.74<br>(0.74) | 2.24 | 0.13 | 1<sub>618.4</sub> |
| Boldness × cluster × stage | 2.63 **<br>(2.63) | 7.96 | **0.004** | 1<sub>620.3</sub> |
| *Random Effects* | *Δ log-lik* | *χ<sup>2</sup>* | *p* | *df* |
| population | 38.0 *** | 16.1 | **<0.001** | 1 |
| region | 0.0 | 0 | 1 | 1 |

|  |  |
| --- | --- |
| $\sigma^2$ | 0.33 |
| ICC | 0.06 |
| Observations | 631 |
| Marginal R<sup>2</sup> | 0.647 |

**Growth rate including activity (mean speed) as predictor**
| *Predictors* | *SumSq*<br>*(mean sq)* | *F-value* | *p* | *df (Num<sub>dem</sub>)* |
| --- | --- | --- | --- | --- |
| Activity | 0.49<br>(0.49) | 1.47 | 0.22 | 1<sub>475.2</sub> |
| cluster | 21.09 ***<br>(21.09) | 62.36 | **<0.001** | 1<sub>166.5</sub> |
| stage | 0.88<br>(0.88) | 2.62 | 0.11 | 1<sub>476.2</sub> |
| Activity × cluster | 0.00<br>(0.00) | 0.00 | 0.98 | 1<sub>475.2</sub> |
| Activity × stage | 0.42<br>(0.42) | 1.25 | 0.27 | 1<sub>474.0</sub> |
| cluster × stage | 0.04<br>(0.04) | 0.11 | 0.74 | 1<sub>476.1</sub> |
| Activity × cluster × stage | 0.07<br>(0.07) | 0.02 | 0.89 | 1<sub>474.0</sub> |
| *Random Effects* | *Δ log-lik* | *χ<sup>2</sup>* | *p* | *df* |
| population | 6.1 *** | 11.5 | **<0.001** | 1 |
| region | 0.0 | 0 | 1 | 1 |

|  |  |
| --- | --- |
| $\sigma^2$ | 0.34 |
| ICC | 0.06 |
| Observations | 489 |
| Marginal R<sup>2</sup> | 0.631 |

**Growth rate including exploration (mean distance to shelter) as predictor**
| *Predictors* | *SumSq*<br>*(mean sq)* | *F-value* | *p* | *df (Num<sub>dem</sub>)* |
| --- | --- | --- | --- | --- |
| Exploration | 0.02<br>(0.02) | 0.07 | 0.80 | 1<sub>474.2</sub> |
| cluster | 77.23 ***<br>(77.23) | 230.52 | **<0.001** | 1<sub>16.5</sub> |
| stage | 1.68*<br>(1.68) | 5.02 | **0.025** | 1<sub>479.1</sub> |
| Exploration × cluster | 0.01<br>(0.01) | 0.04 | 0.84 | 1<sub>474.2</sub> |
| Exploration × stage | 2.12 *<br>(2.12) | 6.34 | **0.01** | 1<sub>475.8</sub> |
| cluster × stage | 0.37<br>(0.37) | 1.10 | 0.29 | 1<sub>479.1</sub> |
| Exploration × cluster × stage | 0.95<br>(0.95) | 2.84 | 0.092 | 1<sub>475.8</sub> |
| *Random Effects* | *Δ log-lik* | *χ<sup>2</sup>* | *p* | *df* |
| population | 7.7 *** | 12.5 | **<0.001** | 1 |
| region | 0.0 | 0 | 1 | 1 |
| $\sigma^2$ | 0.34 | | | |
| ICC | 0.07 |  |  |  |
| Observations | 489 |  |  |  |
| Marginal R<sup>2</sup> | 0.634 |  |  |  |
| * $p < 0.05$ ** $p < 0.01$ *** $p < 0.001$ | | | | |

**Table 5.**
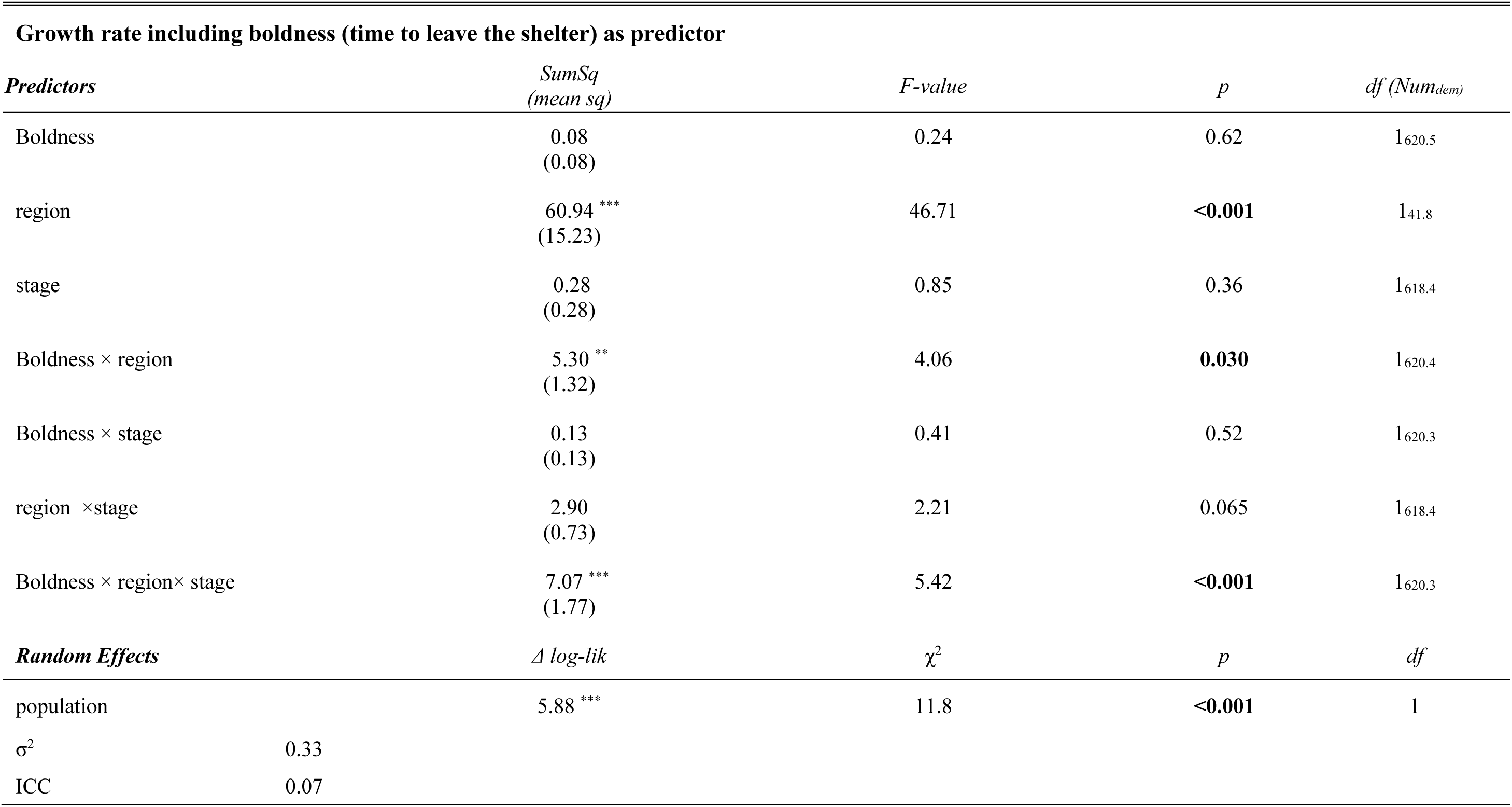
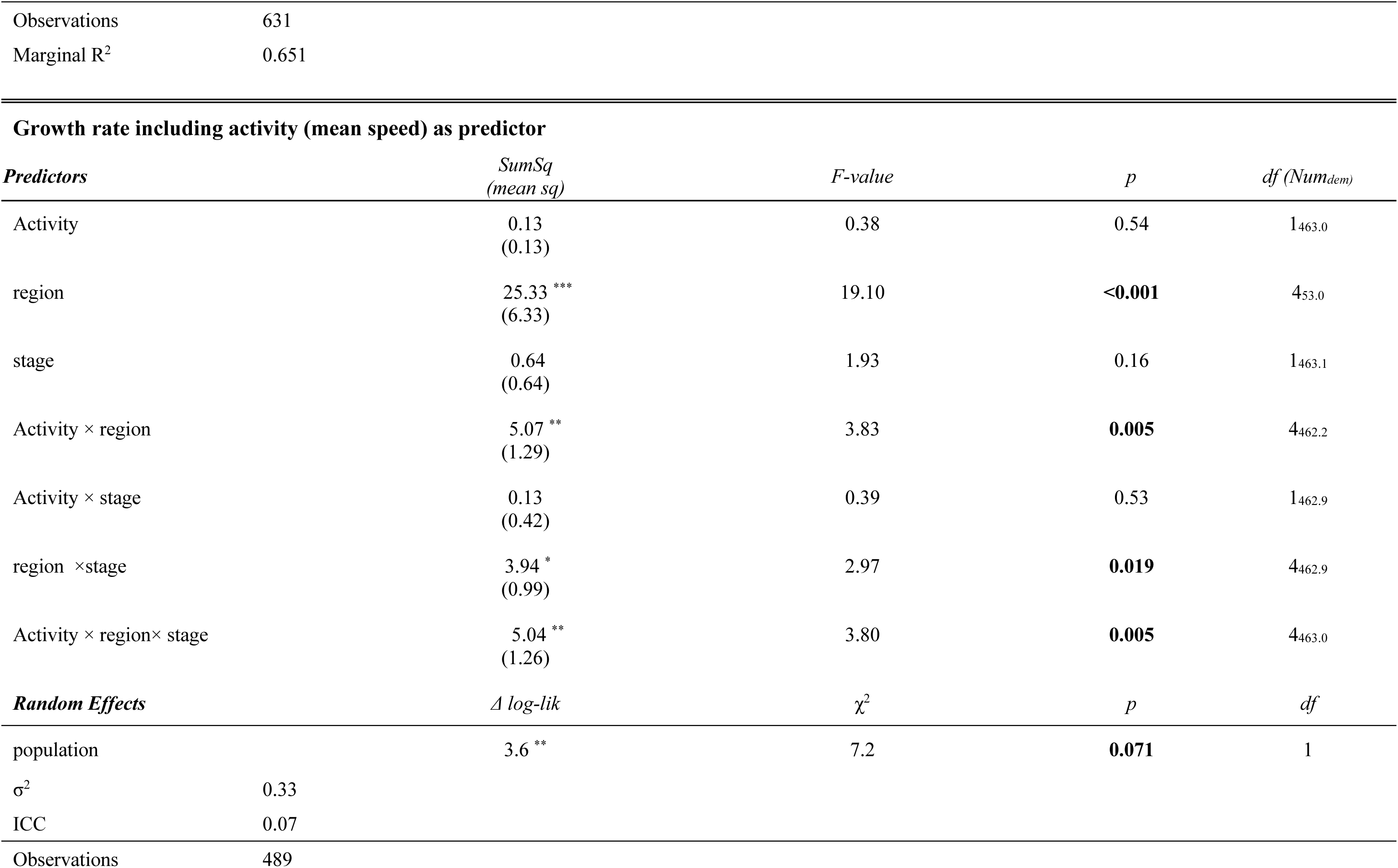
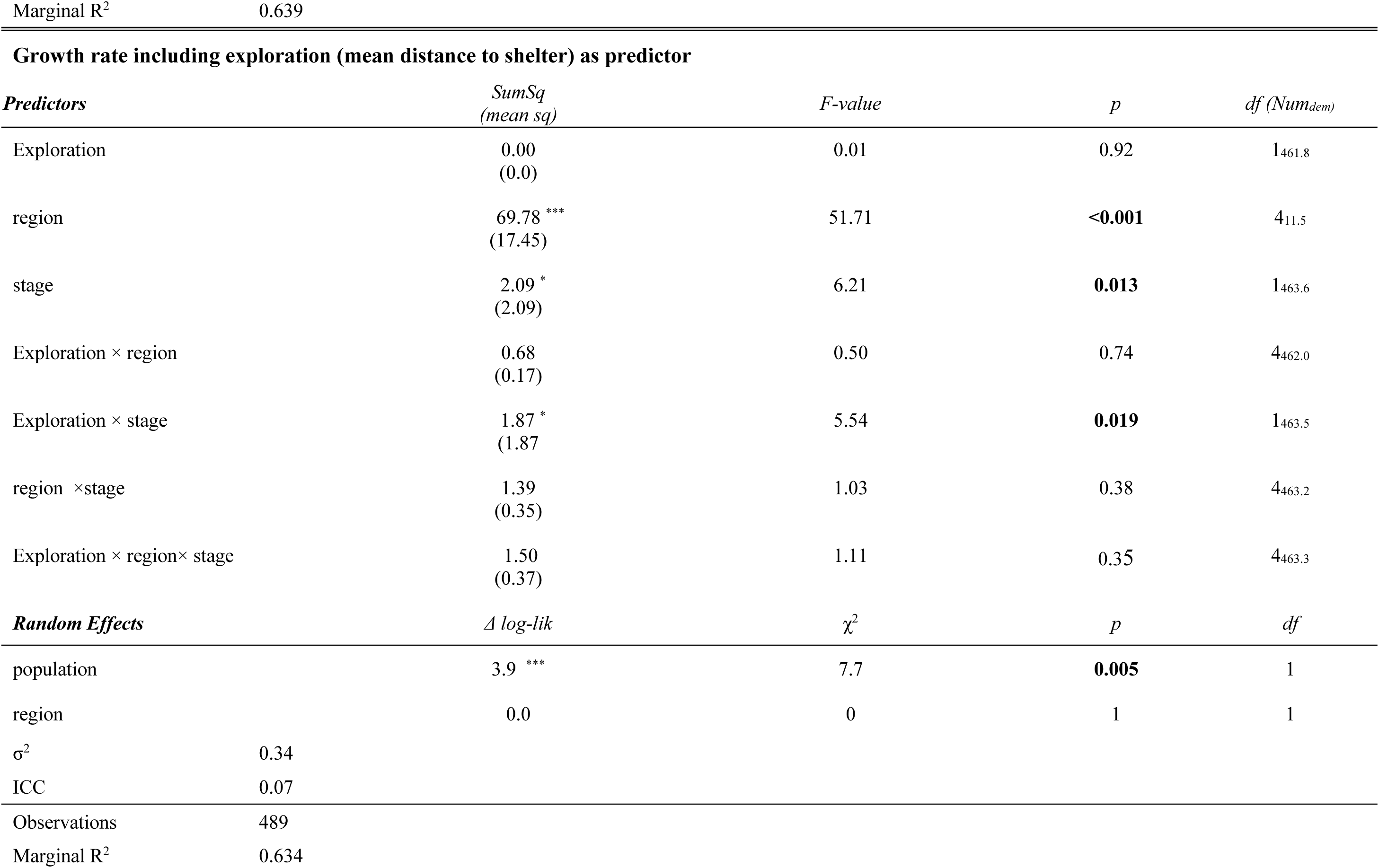
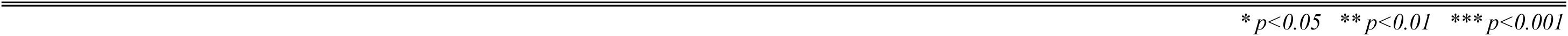
Influence of behavior on life history (growth rate) across regions and developmental stages. Results for statistical analyses evaluating growth rate in moor frog tadpoles and froglets originating from five different regions across a 1700 latitudinal gradient and raised in a common garden experiment (from South to North: Hanover (Germany), and Skåne, Uppsala, Umeå, Luleå (Sweden)). Growth rate models included the behavior of interest, region, developmental stage and their interaction as predictors, and population of origin as a random intercept effect.

#### Growth/activity

Growth rate was not affected by the three-way interaction between activity, genetic cluster and developmental stage, or by any two-way interactions between these factors (Figure 3B, Table 4). Statistical models incorporating the region of origin indicated a significant three-way interaction between activity, region and developmental stage (Fig S2B, Table 5). This effect was driven by the positive association between growth rate and activity in froglets in Uppsala (estimate = 0.73 ± 0.25 *SE*, *P* = 0.032) and Luleå (estimate = 0.89 ± 0.35 *SE*, *P* = 0.059; see table S3). Additionally, it was driven by i) differential associations of growth rate/activity in Hanover-Uppsala contrasts, as well as contrasts of Hanover, Skåne and Umeå against Luleå froglets (see Table S4; Figure S2A); and ii) in contrast with this relationship when measured as froglets in Uppsala (estimate _Froglet_ = -0.01 ± 0.07 *SE*, Tadpole vs Froglet: t = 2.91, *P* = 0.004; Table S5; FigureS2A) and Luleå (estimate _Froglet_ = -0.01 ± 0.07 *SE*, Tadpole vs Froglet: t = 2.91, *P* = 0.004; Table S5; FigureS2A). The random population effect in activity was significant in models incorporating genetic cluster or region (Tables 4 and 5 respectively).

#### Growth/exploration

Growth rate was marginally affected by the three-way interaction between exploration, genetic cluster and developmental stage, and the exploration x developmental stage interaction was significant (Table 4). In the North cluster, there was a negative relationship between exploration and growth rate observed at tadpole, but not at froglet stage (Figure 3C, Table S2). Statistical models incorporating the region of origin indicated an interaction between exploration and developmental stage (Fig S2B, Table 5) driven by a significant change in the direction of this association in Luleå (Tadpole: estimate = -0.15 ± 0.08 *SE*; Froglet: estimate = 0.21 ± 0.15 *SE*; t = -2.12, *P* = 0.034; see Table S5; Figure S2A). The random population effect in activity was significant in models incorporating genetic cluster or region (Tables 4 and 5 respectively).

## DISCUSSION

Our investigation of behavioural patterns in *R. arvalis* at two different developmental stages along an extensive geographical gradient highlights the significant variation in proactivity levels across developmental stages, genetic clusters, and regions in this species. We found no individual repeatability of boldness during tadpole stage, but significant repeatability of this trait during froglet stage. Additionally, our analyses of individual repeatability across metamorphosis showed divergent patterns between behaviours quantified, with activity levels showing significant repeatability that was not observed in exploration and boldness. Our results reveal reversals in behaviour across developmental stages in *R. arvalis*, with all traits quantified showing shifts between the tadpole and froglet stages. Finally, growth rate showed complex interactions with boldness, activity, and exploration, varying by genetic cluster, region, and developmental stage.

### Consistency of inter-individual differences in proactivity and across ontogeny

The repeatability of behaviour was assessed only on the time to emerge from the shelter. The froglets had a repeatable time to emerge from the shelter but not the tadpoles. This suggests that the ontogenetic timing is important to consider in studies investigating the consistency of inter-individual differences in behaviour and assessing personalities. Indeed, studies across taxa support the idea that boldness often develops later in ontogeny, as observed in several vertebrate taxa where boldness may appear only after sexual maturation (Petelle et al. 2013, Polverino et al. 2016, Delval et al. 2020). Moreover, the environmental conditions experienced by previous generations and throughout the lifetime of an individual may also influence the emergence of personalities (e.g. Tariel et al. 2020; Plaskonka et al. 2024). In this study, the results confirm that the time to emerge from a shelter is a proxy of the bold-shy axis of personality in *R. arvalis* and a late emergence of boldness during ontogeny. As it was not the main aim of this study, further assessment of within-stage behavioural consistency is necessary before drawing firm conclusions about when personality emerges and how it is shaped by demographic, geographical and local factors.

Repeatability estimates across life stages indicated no consistency in individuals’ boldness and explorations over ontogeny while activity was significantly repeatable across tadpole and froglet developmental stages. Previous studies across taxa have reported consistency in individuals’ proactivity levels across life stages (reviewed by Koenig and Ousterhout 2018). In anurans, individual consistency in proactivity levels before and after metamorphosis has been observed in the spotted salamander (Koenig and Ousterhout 2018), and in the activity and exploration levels of *Pelophylax ridibundus* from a single population from northern Germany (Wilson and Krause, 2012). However, neither Plaskonka et al. (2024) in a single population of Polish *R. arvalis* or Brodin et al. (2013) in several northern Swedish populations of *R. temporaria* found significant repeatability across life stages. Our results are in broad agreement with the previous studies, suggesting that repeatability across life stages can be weak and depend on the environmental constraints. Indeed, our southernmost population exhibited the lowest changes in proactivity traits across life stages, while there was a tendency to gradually increase in change across life stages with increasing latitude (see next parts of discussion). Together, these findings suggest that individual consistency in behaviour across life stages may vary across taxa due to the adaptive value of consistent behavioural responses, shaped to a larger extent by ecological factors than by underlying genetic drivers of behavioural syndromes.

### Proactive behaviours of tadpoles along the latitudinal gradient

Activity was higher in tadpoles from the North genetic cluster and in the three northernmost regions compared to the South genetic cluster and the two southernmost regions, respectively. In addition, tadpoles from the two northernmost regions also showed a higher activity than the latitudinally intermediate Uppsala region. Consistently, tadpoles from the northernmost region (Luleå) exhibited higher boldness than those from other regions and tadpoles from the most southernmost region (Hanover) showed lower boldness than those from other regions, including regions within the same genetic cluster. Altogether, these results suggest that both genetic clusters, resulting from post-glacial colonization events and subsequent demographic processes, and contemporary selection at the regional scale shaped the proactive behaviour of tadpoles. This is in line with our previous studies showing a complex interplay between effects of historical and contemporary processes on the genetic structure (Cortazar Chinarro et al., 2017; 2018, Meyer-Lucht et al., 2019, Rödin-Mörch et al., 2019) and larval life history traits (Luquet et al., 2019) of *R. arvalis* populations along the latitudinal gradient. These findings likewise align with the broader framework of POLS theory, which proposes that fast-slow life history strategies correlate with personality traits along environmental gradients (Reale et al. 2010). Our results on *R. arvalis* tadpoles provide empirical support for this concept, particularly in the context of ectotherms facing strong seasonal time constraints at early developmental stages, as previously observed in anti-predator responses across lizard species (Samia et al. 2015). However, an interplay on proactive behaviour of *R. arvalis* tadpoles is difficult to disentangle and seems also depend on the behavioural trait considered and regional environmental conditions. Indeed, activity of tadpoles is strongly influenced at the cluster scale with low variation within genetic cluster. In contrast, tadpole boldness showed strong differences among regions within and between genetic clusters, resulting in no difference in tadpole boldness between the genetic clusters. Tadpole exploration was not influenced by latitude neither at genetic cluster nor at regional levels. This suggests that boldness variation along the latitudinal gradient results from contemporary selection due to environmental constraints for bolder genotypes in the northern populations, while activity variation along the gradient results from the genetic divergence due to historical post-recolonization. Findings of trait-specific behavioural responses in our latitudinal gradient study further contribute to revisitations of POLS theory, demonstrating that different personality traits can evolve independently in response to the influence of both historical events and ongoing environmental pressures (Dingemanse et al., 2010; Wolf and Weissing, 2012, Royauté et al. 2018).

### Decoupling of proactive behaviours across ontogeny along the latitudinal gradient

Interestingly, the proactive behaviour of froglets along the gradient is almost the reverse of the tadpole behaviour. Froglets from the North cluster exhibited reduced boldness and exploration as compared to the South, and froglets from the two northernmost regions were shyer and explored less than froglets from other regions. There was no change in froglet activity along the gradient. This result aligns well with the extended POLS theory, predicting that behaviours on the proactive-reactive continuum can be decoupled across ontogeny (Montiglio et al., 2018). The observed decoupling is likely due to fluctuating selective pressures that amphibians face as they transition from aquatic to terrestrial environments following metamorphosis (Van Buskirk, 2002; Relyea and Hoverman, 2003, Orizaola and Braña, 2005). Our results suggest that in high latitude populations, tadpoles may benefit from increased boldness and activity in aquatic environments, where rapid growth and resource acquisition are critical for survival as the tadpoles have to metamorphose before the onset of winter (Laugen et al., 2003, Luquet et al., 2019, Orizaola et al., 2013). However, once living in terrestrial environments, the cost of being overly bold or explorative in harsher northern climates could outweigh the benefits. Specifically, in high-latitude populations time for foraging after metamorphosis may be very short due to approaching winter which may select for lower boldness and exploration activity. Indeed, the highest exploration level was found in the Hanover population which experiences the longest season length along the gradient, while Luleå experiences the shortest, suggesting that this trait may be related to the time available for resource acquisition after metamorphosis. In juvenile *R. arvalis* terrestrial growth rates in laboratory are lower in the northern than in the southern populations (Chondrelli, 2024), and studies on growth and survival patterns in Scandinavian *Rana* populations suggest that survival is higher but maturation occurs later in the northernmost populations (Hjernquist et al., 2012; Räsänen et al., 2008; Söderman, 2006). These results suggest that time constraints for terrestrial growth and development may be relaxed along the latitudinal gradient as compared to the more stringent constraints faced by tadpoles, possibly allowing for decoupling of behaviours related to resource acquisition (Sih et al., 2004). A recent study also suggests that the ontogenetic experience of predator presence led to personality emergence in *R. arvalis* tadpoles and that this pattern was lost over metamorphosis (Plaskonka et al. 2024); as predation pressure is known to decrease with latitude (Laurila et al., 2008; Schemske et al., 2009) and could also influence the decoupling of behaviours. Such ontogenetic decoupling of behaviours has been observed in other species, where selective pressures shift as individuals transition between ecological niches (Wuerz and Kruger, 2015, Katsis et al. 2023, Class and Brommer, 2015, Bosco et al., 2017). Thus, our findings underscore the importance of considering both life-history stage and the ecological context when examining behavioural traits and personalities across geographical ranges.

### Relationship between proactive behaviours and larval growth along the latitudinal gradient

Developmental growth is an important life history trait in larval amphibians, and several studies have shown that this trait readily diverges along climatic gradients (e.g., Berven et al. 1983: Laurila et al. 2008, Orizaola et al. 2010). In our previous study focusing on life-history traits, we observed notable differences in larval growth among *R. arvalis* populations deriving from the two post-glaciation colonization routes, with populations from the northern cluster showing a higher growth rate (Luquet et al. 2019; see Figure 3). Behavioural variation of tadpoles observed in this study (mainly higher activity in the North cluster and the northernmost regions, and higher boldness in the northernmost region) appears to globally align with expectations that proactive behaviours coevolve with faster larval growth. However, detailed relationships between proactive behaviours and larval growth showed a strong influence of both genetic cluster and developmental stage. Indeed, larval growth is positively associated with boldness in tadpoles from the South cluster but not in tadpoles from the North one, while in froglets larval growth is positively associated with boldness in the North cluster but not in the South one. At the regional scale, our findings further disentangle the role of historical processes and local adaptation in shaping growth-behaviour relationships. Positive associations between tadpole boldness and larval growth were observed in Uppsala and Skåne, as well as between froglet activity and larval growth in Uppsala. However, these associations were absent in other regions, suggesting that local environmental conditions have more strength in modulating such associations, as previously observed in insects (Golab et al. 2022).

As mentioned above, these different relationships across genetic clusters and among regions are likely driven by both colonization history and climate-induced time constraints, which have together shaped crucial shifts in metabolism and life-history traits (Laurila et al. 2008; Orizaola et al. 2013, Caruso et al. 2020). Additionally, ecological factors such as predation pressure and susceptibility to parasitic infections likely play crucial roles in shaping contrasting relationships between life history and behaviour in different geographical areas. For instance, as predation pressures on amphibians is typically reduced, and parasite diversity tends to decrease with increasing latitude (Roslin et al. 2017, Rhode 1999), the contrasting relationships between growth and proactivity across clusters may reflect the interplay of these ecological pressures. In the North cluster, where both predation and parasitism are reduced, proactive behaviours may provide an advantage later in development, explaining the positive association between boldness and larval growth. Conversely, in the South cluster, where predation and parasite diversity are higher, selection might favour proactivity earlier during tadpole stage, enabling faster growth and providing an adaptive strategy that compensate higher exposure to predators and infection. This divergence in selective pressures likely explains the inconsistencies observed between the genetic clusters, with local adaptation influencing how behaviour and life-history traits co-vary across ontogeny and different regions along the gradient.

### Conclusions

*R. arvalis* populations along a broad latitudinal gradient in northern Europe exhibit clear differences in proactivity levels, showing reversed behavioural patterns before and after metamorphosis. Our findings suggest that natural selective pressures acting on behaviours along the proactive-reactive continuum and life history traits operate differently across developmental stages in this species. These results emphasize the importance of latitudinal gradient studies to elucidate evolutionary processes driving phenotypic variation in behaviour among natural populations. Future research should investigate the role of temperature, predator diversity, and resource availability in shaping these behaviours. Furthermore, linking behavioural variation across developmental stages to specific genomic polymorphisms could provide valuable insights into evolutionary processes, historical selection and the maintenance of behavioural diversity across regions.

## Acknowledgements

We thank Emma Dahl and Soraya Rouifed for help in the field and conducting behavioural assays, and Wouter van der Bijl for help with tracking analyses and visualization. The eggs were collected with permits from the Authority for Nature Protection of Lower Saxony and the Hanover region and respective county boards in Sweden. We thank Anders Hallengren (Länstyrelsen Skåne), Anders Forsgren and Stefan Andersson (Piteå kommun), Thomas Brandt (Ökologische Schutzstation Steinhuder Meer e.V), and Bernd Rittberg and Andreas Jacob (Authority for Nature Protection of Lower Saxony and the Hanover region) for helping locating the sampling locations. The experiment was conducted with a permit (C40/14) from the Ethical Committee for Animal Experiments in Uppsala County.

## Author contributions

M.C-C.; writing original draft, writing-review and editing, A.C-L.; data curation, formal analyses, writing original draft writing-review and editing, E.L.; conceptualization, field work, lab work, writing-review and editing, A.L.; conceptualization, funding, writing-review and editing.

## References

Alford, R. A. (1999). Ecology: resource use, competition, and predation. Tadpoles: the biology of anuran larvae, 240–278.

Amat, I., Desouhant, E., Gomes, E., Moreau, J., Perhaps we should mention shy-bold dichotomy/continuum as a key prediction here?

Monceau, K. (2018). Insect personality: what can we learn from metamorphosis? Current opinion in insect science, 27, 46–51.

Bakker TC (1986). Aggressiveness in sticklebacks (Gasterosteus aculeatus L.): a behaviour-genetic study. Behaviour 98(1-4): 1–144.

Bates, D., Maechler, M., Bolker, B., Walker, S., Christensen, R. H. B., Singmann, H., … & Bolker, M. B. (2015). Package ‘lme4’. convergence, 12(1), 2.

Bégué, L., Tschirren, N., Peignier, M., Szabo, B., & Ringler, E. (2024). Behavioural consistency across metamorphosis in a neotropical poison frog. Evolutionary Ecology, 38(1), 157–174.

Biro, P. A., & Stamps, J. A. (2008). Are animal personality traits linked to life-history productivity? Trends in ecology & evolution, 23(7), 361–368.

Bosco, J. M., Riechert, S. E., & O’Meara, B. C. (2017). The ontogeny of personality traits in the desert funnel-web spider, Agelenopsis lisa (Araneae: Agelenidae). Ethology, 123(9), 648–658.

Bouchard TJ, Loehlin JC (2001). Genes, evolution, and personality. Behavior genetics 31(3): 243–273.

Branson K, Robie AA, Bender J, Perona P, Dickinson MH (2009). High-throughput ethomics in large groups of Drosophila. Nature methods 6(6): 451–457.

Brodin, T. (2009). Behavioral syndrome over the boundaries of life—carryovers from larvae to adult damselfly. Behavioral Ecology, 20(1), 30–37.

Brodin, T., Lind, M. I., Wiberg, M. K., & Johansson, F. (2013). Personality trait differences between mainland and island populations in the common frog (Rana temporaria). Behavioral Ecology and Sociobiology, 67, 135–143.

Cabrera, D., Nilsson, J. R., & Griffen, B. D. (2021). The development of animal personality across ontogeny: a cross-species review. Animal Behaviour, 173, 137–144.

Careau, V., Thomas, D., Humphries, M. M., & Réale, D. (2008). Energy metabolism and animal personality. Oikos, 117(5), 641–653.

Cassidy C, Grange LJ, Garcia C, Bolam SG, Godbold JA (2020). Species interactions and environmental context affect intraspecific behavioural trait variation and ecosystem function. Proceedings of the Royal Society B 287(1919): 20192143.

Chang, C. C., Moiron, M., Sánchez-Tójar, A., Niemelä, P. T., & Laskowski, K. L. (2024). What is the meta-analytic evidence for life-history trade-offs at the genetic level? Ecology Letters, 27(1), e14354.

Chondrelli, N. (2024). Thermal mismatch in high latitude host-parasite interactions (Doctoral dissertation, Acta Universitatis Upsaliensis).

Class, B., & Brommer, J. E. (2015). A strong genetic correlation underlying a behavioural syndrome disappears during development because of genotype–age interactions. Proceedings of the Royal Society B: Biological Sciences, 282(1809), 20142777.

Cohen J (1985). Metamorphosis: introduction, usages, and evolution. Metamorphosis Oxford University Press, Oxford, UK: 1–19.

Cortázar-Chinarro M, Lattenkamp EZ, Meyer-Lucht Y, Luquet E, Laurila A, Höglund J (2017). Drift, selection, or migration? Processes affecting genetic differentiation and variation along a latitudinal gradient in an amphibian. BMC Evolutionary Biology 17(1): 1–14.

Cortazar-Chinarro, M., Meyer-Lucht, Y., Laurila, A., & Höglund, J. (2018). Signatures of historical selection on MHC reveal different selection patterns in the moor frog (Rana arvalis). Immunogenetics, 70, 477–484.

Culumber, Z. W. (2022). Variation in behavioral traits across a broad latitudinal gradient in a livebearing fish. Evolutionary Ecology, 36(1), 75–91.

Delval, I., Fernández-Bolaños, M., & Izar, P. (2020). A longitudinal assessment of behavioral development in wild capuchins: Personality is not established in the first 3 years. American Journal of Primatology, 82(11), e23116.

Dingemanse, N. J., & Dochtermann, N. A. (2013). Quantifying individual variation in behaviour: mixed-effect modelling approaches. Journal of Animal Ecology, 82(1), 39–54.

Fischer AG (1960). Latitudinal variations in organic diversity. Evolution 14(1): 64–81.

Foster SA, Wund MA, Baker JA (2015). Evolutionary influences of plastic behavioral responses upon environmental challenges in an adaptive radiation. Integrative and Comparative Biology 55(3): 406–417.

Gerlai R, Csányi V (1990). Genotype-environment interaction and the correlation structure of behavioral elements in paradise fish (Macropodus opercularis). Physiology & behavior 47(2): 343–356.

Gopal, A. C., Alujević, K., & Logan, M. L. (2023). Temperature and the pace of life. Behavioral Ecology and Sociobiology, 77(5), 59.

Golab, M. J., Sniegula, S., & Brodin, T. (2022). Cross-latitude behavioural axis in an adult damselfly Calopteryx splendens (Harris, 1780). Insects, 13(4), 342.

Gosner KL (1960). A simplified table for staging anuran embryos and larvae with notes on identification. Herpetologica 16(3): 183–190.

Groothuis TG, Carere C (2005). Avian personalities: characterization and epigenesis. Neuroscience & Biobehavioral Reviews 29(1): 137–150.

Halekoh U, Højsgaard S (2014). “A Kenward-Roger Approximation and Parametric Bootstrap Methods for Tests in Linear Mixed Models – The R Package pbkrtest.” Journal of Statistical Software, 59(9), 1–30.

Healy, K., Ezard, T. H., Jones, O. R., Salguero-Gómez, R., & Buckley, Y. M. (2019). Animal life history is shaped by the pace of life and the distribution of age-specific mortality and reproduction. Nature Ecology & Evolution, 3(8), 1217–1224.

Higgins TA, Wilcox RC, Germain RR, Tarwater CE (2022). Behavioral traits vary with intrinsic factors and impact local survival in Song Sparrows (*Melospiza melodia*). The Wilson Journal of Ornithology.

Katsis, A. C., Common, L. K., Hauber, M. E., Colombelli-Négrel, D., & Kleindorfer, S. (2023). From nestling to adult: personality traits are consistent within but not across life stages in a wild songbird. Behaviour, 160(8-9), 701–734.

Kelleher SR, Silla AJ, Byrne PG (2018). Animal personality and behavioral syndromes in amphibians: a review of the evidence, experimental approaches, and implications for conservation. Behavioral Ecology and Sociobiology 72(5): 1–26.

Knopp T, Merilä J (2009). The postglacial recolonization of Northern Europe by Rana arvalis as revealed by microsatellite and mitochondrial DNA analyses. Heredity 102(2): 174–181.

Koenig, A. M., & Ousterhout, B. H. (2018). Behavioral syndrome persists over metamorphosis in a pond-breeding amphibian. Behavioral Ecology and Sociobiology, 72, 1–12.

Kuznetsova A, Brockhoff PB, Christensen RHB (2017). “lmerTest Package: Tests in Linear Mixed Effects Models.” Journal of Statistical Software, 82(13), 1–26.

Laurila, A., Lindgren, B., & Laugen, A. T. (2008). Antipredator defenses along a latitudinal gradient in Rana temporaria. Ecology, 89(5), 1399–1413.

Laurila, A., Pakkasmaa, S. & Merilä, J, (2006) Population divergence in growth rate and antipredator defenses in *Rana arvalis*. Oecologia, 147, 585–595.

Lenth, R., & Lenth, M. R. (2018). Package ‘lsmeans’. The American Statistician, 34(4), 216–221.

Luquet E, Rödin Mörch P, Cortázar-Chinarro M, Meyer-Lucht Y, Höglund J, Laurila A (2019). Post-glacial colonization routes coincide with a life-history breakpoint along a latitudinal gradient. Journal of Evolutionary Biology 32(4): 356–368.

Luquet, E., Léna, J. P., Miaud, C., & Plénet, S. (2015). Phenotypic divergence of the common toad (Bufo bufo) along an altitudinal gradient: evidence for local adaptation. Heredity, 114(1), 69–79.

Mazué GP, Dechaume-Moncharmont F-X, Godin J-GJ (2015). Boldness–exploration behavioral syndrome: interfamily variability and repeatability of personality traits in the young of the convict cichlid (Amatitlania siquia). Behavioral Ecology 26(3): 900–908.

Meyer-Lucht Y, Luquet E, Jóhannesdóttir F, Rödin-Mörch P, Quintela M, Richter-Boix A, et al (2019). Genetic basis of amphibian larval development along a latitudinal gradient: Gene diversity, selection and links with phenotypic variation in transcription factor C/EBP-1. Molecular Ecology 28(11): 2786–2801.

Mitchell D, Beatty ET, Cox PK (1977). Behavioral differences between two populations of wild rats: implications for domestication research. Behavioral Biology 19(2): 206–216.

Montiglio, P. O., Dammhahn, M., Messier, G. D. & Réale, D. The pace-of-life syndrome revisited: the role of ecological conditions and natural history on the slow-fast continuum. Behavioral Ecology and Sociobiology 72, 116 (2018).

Nakagawa, S., & Schielzeth, H. (2010). Repeatability for Gaussian and non-Gaussian data: a practical guide for biologists. Biological Reviews, 85(4), 935–956.

Orizaola, G., & Brana, F. (2005). Plasticity in newt metamorphosis: the effect of predation at embryonic and larval stages. Freshwater Biology, 50(3), 438–446.

Orizaola, G., Dahl, E., Nicieza, A. G., & Laurila, A. (2013). Larval life history and anti-predator strategies are affected by breeding phenology in an amphibian. Oecologia, 171, 873–881.

Orizaola, G., Quintela, M., & Laurila, A. (2010). Climatic adaptation in an isolated and genetically impoverished amphibian population. Ecography, 33(4), 730–737.

Petelle, M. B., McCoy, D. E., Alejandro, V., Martin, J. G., & Blumstein, D. T. (2013). Development of boldness and docility in yellow-bellied marmots. Animal Behaviour, 86(6), 1147–1154.

Płaskonka, B., Zaborowska, A., Mikulski, A., & Pietrzak, B. (2024). Predation Risk Experienced by Tadpoles Shapes Personalities Before but Not After Metamorphosis. Ecology and Evolution, 14(11), e70532.

Polverino, G., Cigliano, C., Nakayama, S., & Mehner, T. (2016). Emergence and development of personality over the ontogeny of fish in absence of environmental stress factors. Behavioral Ecology and Sociobiology, 70, 2027–2037.

Poulin, R., & Leung, T. L. F. (2011). Latitudinal gradient in the taxonomic composition of parasite communities. Journal of Helminthology, 85(3), 228–233.

R Core Team (2024). _R: A Language and Environment for Statistica Computing. R Foundation for Statistical Computing, Vienna, Austria. https://www.R-project.org/.

Räsänen, K., Söderman, F., Laurila, A., & Merilä, J. (2008). Geographic variation in maternal investment: acidity affects egg size and fecundity in Rana arvalis. Ecology, 89(9), 2553–2562.

Réale, D. et al. Personality and the emergence of the pace-of-life syndrome concept at the population level. Philosophical Transactions of the Royal Society B: Biological Sciences 365, 4051–4063 (2010).

Relyea R.A. & Hoverman J.T. (2003) The impact of larval predators and competitors on the morphology and fitness of juvenile treefrogs. Oecologia, 134, 596 – 604.

Rödin-Mörch P, Luquet E, Meyer-Lucht Y, Richter-Boix A, Höglund J, Laurila A (2019). Latitudinal divergence in a widespread amphibian: Contrasting patterns of neutral and adaptive genomic variation. Molecular Ecology 28(12): 2996–3011.

Roslin, T., Hardwick, B., Novotny, V., Petry, W. K., Andrew, N. R., Asmus, A., … & Slade, E. M. (2017). Higher predation risk for insect prey at low latitudes and elevations. Science, 356(6339), 742–744.

Rowe L, Ludwig D (1991). Size and timing of metamorphosis in complex life cycles: time constraints and variation. Ecology 72(2): 413–427.

Samia, D. S., Møller, A. P., Blumstein, D. T., Stankowich, T., & Cooper Jr, W. E. (2015). Sex differences in lizard escape decisions vary with latitude, but not sexual dimorphism. Proceedings of the Royal Society B, 282(1805), 20150050.

Schemske, D. W., Mittelbach, G. G., Cornell, H. V., Sobel, J. M., & Roy, K. (2009). Is there a latitudinal gradient in the importance of biotic interactions?. Annu. Rev. Ecol. Evol. Syst., 40, 245–269.

Sih, A., Bell, A. M., Johnson, J. C., & Ziemba, R. E. (2004). Behavioral syndromes: an integrative overview. The Quarterly Review of Biology, 79(3), 241–277.

Sillero N, Campos J, Bonardi A, Corti C, Creemers R, Crochet P-A et al (2014). Updated distribution and biogeography of amphibians and reptiles of Europe. Amphibia-Reptilia 35(1): 1–31.

Söderman, F. (2006). Comparative population ecology in moor frogs with particular reference to acidity (Doctoral dissertation, Acta Universitatis Upsaliensis).

Spiegel, O., Leu, S. T., Sih, A., Godfrey, S. S., & Bull, C. M. (2015). When the going gets tough: behavioural type-dependent space use in the sleepy lizard changes as the season dries. Proceedings of the Royal Society B: Biological Sciences, 282(1819), 20151768.

Stamps, J. A. (2007). Growth-mortality tradeoffs and ‘personality traits’ in animals. Ecology letters, 10(5), 355–363.

Stearns SC (1992) The evolution of life histories. Oxford University Press, Oxford New York

Stoffel, M. A., Nakagawa, S., & Schielzeth, H. (2017). rptR: Repeatability estimation and variance decomposition by generalized linear mixed-effects models. Methods in Ecology and Evolution, 8(11), 1639–1644.

Tariel, J., Plénet, S., & Luquet, E. (2020). How do developmental and parental exposures to predation affect personality and immediate behavioural plasticity in the snail Physa acuta? Proceedings of the Royal Society B, 287(1941), 20201761.

Therneau, T. M., & Lumley, T. (2015). Package ‘survival’. R Top Doc, 128(10), 28–33.

Van Buskirk J. (2002) Phenotypic lability and the evolution of predator-induced plasticity in tadpoles. Evolution, 56, 361 – 370.

Villegas-Ríos, D., Réale, D., Freitas, C., Moland, E., & Olsen, E. M. (2018). Personalities influence spatial responses to environmental fluctuations in wild fish. Journal of Animal Ecology, 87(5), 1309–1319.

Wilbur HM (1980). Complex life cycles. Annual review of Ecology and Systematics 11: 67–93.

Willig, M. R., Kaufman, D. M., & Stevens, R. D. (2003). Latitudinal gradients of biodiversity: pattern, process, scale, and synthesis. Annual review of ecology, evolution, and systematics, 34(1), 273–309.

Wilson AD, Krause J (2012). Personality and metamorphosis: is behavioral variation consistent across ontogenetic niche shifts? Behavioral Ecology 23(6): 1316–1323.

Wilson DS, Clark AB, Coleman K, Dearstyne T (1994). Shyness and boldness in humans and other animals. Trends in ecology & evolution 9(11): 442–446.

Wuerz, Y., & Krüger, O. (2015). Personality over ontogeny in zebra finches: long-term repeatable traits but unstable behavioural syndromes. Frontiers in Zoology, 12(Suppl 1), S9.

