## SupplementaryTables1-5&Figures1-2 for "Metamorphosis reverses the behavioural phenotype in *Rana arvalis* along a latitudinal gradient"

**Table S1.** Independent contrasts of behavior models by developmental stage.

**Table S2.** Independent contrasts of growth rate models by genetic cluster or by developmental stage.

**Table S3.** Trends of growth rate and behavior association by region and developmental stage.

**Table S4.** Independent contrasts of growth rate models by developmental stage.

**Table S5.** Independent contrasts of growth rate models by region.

**Figure S1.** Consistency of individual behaviour across developmental stages.

**Figure S2.** Association between behaviour and life history in moor frogs by region.

**Table S1. Independent contrasts of behavior models by developmental stage.** Statistical contrasts between regions for significant interaction effects in models evaluating behavioral traits across tadpole and froglet stages in moor frogs originated from 5 different regions across a 1700 latitudinal gradient and raised in a common garden experiment. Contrasts were computed on estimated marginal means. Significant p-values adjusted for false discovery rate in bold.

**Boldness:** Time to leave the shelter

| *Dev.stage* | *contrast* | *ratio* | *SE* | *df* | *z.ratio* | *p* |
|---|---|---|---|---|---|---|
| Tadpole | Hanover - Skåne | 0.58 | 0.13 | Inf | -2.40 | **0.040** |
| | Hanover - Uppsala | 0.69 | 0.15 | Inf | -1.70 | 0.14 |
| | Hanover - Umeå | 0.74 | 0.18 | Inf | -1.26 | 0.29 |
| | Hanover - Luleå | 0.38 | 0.09 | Inf | -4.05 | **<0.001** |
| | Skåne - Uppsala | 1.17 | 0.19 | Inf | 0.98 | 0.36 |
| | Skåne - Umeå | 1.25 | 0.25 | Inf | 1.14 | 0.32 |
| | Skåne - Luleå | 0.65 | 0.12 | Inf | -2.32 | **0.040** |
| | Uppsala - Umeå | 1.07 | 0.19 | Inf | 0.38 | 0.70 |
| | Uppsala - Luleå | 0.55 | 0.09 | Inf | -3.45 | **0.001** |
| | Umeå - Luleå | 0.51 | 0.08 | Inf | -4.15 | **<0.001** |
| Froglet | Hanover - Skåne | 1.27 | 0.34 | Inf | 0.88 | 0.42 |
| | Hanover - Uppsala | 0.98 | 0.26 | Inf | -0.07 | 0.94 |
| | Hanover - Umeå | 2.01 | 0.54 | Inf | 2.58 | **0.019** |
| | Hanover - Luleå | 2.47 | 0.67 | Inf | 3.31 | **0.002** |
| | Skåne - Uppsala | 0.77 | 0.14 | Inf | -1.36 | 0.25 |
| | Skåne - Umeå | 1.59 | 0.31 | Inf | 2.38 | **0.028** |
| | Skåne - Luleå | 1.95 | 0.38 | Inf | 3.40 | **0.002** |
| | Uppsala - Umeå | 2.05 | 0.37 | Inf | 3.94 | **<0.001** |
| | Uppsala - Luleå | 2.51 | 0.47 | Inf | 4.91 | **<0.001** |
| | Umeå - Luleå | 1.22 | 0.23 | Inf | 1.07 | 0.35 |

**Activity:** mean speed

| *Dev.stage* | *contrast* | *estimate* | *SE* | *df* | *t-value* | *p* |
|---|---|---|---|---|---|---|
| Tadpole | Hanover - Skåne | 0.03 | 0.07 | 9.33 | 0.50 | 0.69 |
| | Hanover - Uppsala | -0.11 | 0.07 | 9.47 | -1.71 | 0.15 |

| | | | | | | |
|---|---|---|---|---|---|---|
| | Hanover - Umeå | -0.21 | 0.07 | 11.01 | -3.09 | **0.022** |
| | Hanover - Luleå | -0.22 | 0.07 | 10.35 | -3.24 | **0.022** |
| | Skåne - Uppsala | -0.15 | 0.05 | 9.54 | -3.13 | **0.022** |
| | Skåne - Umeå | -0.25 | 0.05 | 12.72 | -4.85 | **0.002** |
| | Skåne - Luleå | -0.26 | 0.05 | 11.33 | -5.16 | **0.002** |
| | Uppsala - Umeå | -0.10 | 0.05 | 11.01 | -2.03 | 0.095 |
| | Uppsala - Luleå | -0.11 | 0.05 | 10.03 | -2.23 | 0.081 |
| | Umeå - Luleå | -0.00 | 0.05 | 9.50 | -0.15 | 0.87 |
| Froglet | Hanover - Skåne | 0.01 | 0.09 | 29.03 | 0.13 | 0.90 |
| | Hanover - Uppsala | -0.05 | 0.09 | 26.55 | -0.53 | 0.78 |
| | Hanover - Umeå | 0.04 | 0.09 | 32.00 | 0.40 | 0.78 |
| | Hanover - Luleå | 0.09 | 0.10 | 44.57 | 0.92 | 0.73 |
| | Skåne - Uppsala | -0.06 | 0.06 | 19.37 | -1.02 | 0.73 |
| | Skåne - Umeå | 0.03 | 0.06 | 29.64 | 0.40 | 0.78 |
| | Skåne - Luleå | 0.08 | 0.07 | 55.29 | 1.06 | 0.73 |
| | Uppsala - Umeå | 0.08 | 0.06 | 22.89 | 1.40 | 0.73 |
| | Uppsala - Luleå | 0.14 | 0.07 | 47.43 | 1.92 | 0.61 |
| | Umeå - Luleå | 0.06 | 0.08 | 60.42 | 0.71 | 0.77 |

**Exploration:** mean distance to shelter

| ***Dev.stage*** | ***contrast*** | ***estimate*** | ***SE*** | ***df*** | ***t-value*** | ***p*** |
|---|---|---|---|---|---|---|
| Tadpole | Hanover - Skåne | -2.40 | 2.49 | 12.88 | -0.96 | 0.507 |
| | Hanover - Uppsala | 2.60 | 2.50 | 12.95 | 1.03 | 0.507 |
| | Hanover - Umeå | -0.62 | 2.52 | 13.33 | -0.24 | 0.836 |
| | Hanover - Luleå | 0.52 | 2.49 | 12.81 | 0.21 | 0.836 |
| | Skåne - Uppsala | 5.00 | 1.78 | 13.23 | 2.80 | 0.146 |
| | Skåne - Umeå | 1.77 | 1.80 | 14.01 | 0.98 | 0.507 |
| | Skåne - Luleå | 2.92 | 1.77 | 12.94 | 1.65 | 0.407 |
| | Uppsala - Umeå | -3.22 | 1.81 | 14.16 | -1.78 | 0.407 |
| | Uppsala - Luleå | -2.07 | 1.77 | 13.09 | -1.16 | 0.507 |
| | Umeå - Luleå | 1.15 | 1.80 | 13.86 | 0.64 | 0.664 |

| | | | | | | |
|---|---|---|---|---|---|---|
| Froglet | Hanover - Skåne | 15.86 | 4.40 | 104.87 | 3.60 | **0.001** |
| | Hanover - Uppsala | 12.06 | 4.25 | 94.08 | 2.83 | **0.011** |
| | Hanover - Umeå | 18.33 | 4.58 | 118.61 | 4.00 | **0.001** |
| | Hanover - Luleå | 24.97 | 5.06 | 153.78 | 4.92 | **<0.001** |
| | Skåne - Uppsala | -3.79 | 2.51 | 48.77 | -1.51 | 0.153 |
| | Skåne - Umeå | 2.47 | 3.03 | 93.08 | 0.81 | 0.417 |
| | Skåne - Luleå | 9.11 | 3.78 | 161.41 | 2.44 | **0.026** |
| | Uppsala - Umeå | 6.27 | 2.81 | 72.59 | 2.22 | **0.041** |
| | Uppsala - Luleå | 12.91 | 3.55 | 141.14 | 3.63 | **0.001** |
| | Umeå - Luleå | 6.638 | 3.938 | 186.173 | 1.686 | 0.117 |

**Table S2. Independent contrasts of growth rate models by genetic cluster or by developmental stage.** Statistical contrasts of significant interaction effects between clusters or developmental stages in models evaluating growth rate in moor frog tadpoles and froglets divided in two genetic clusters based on their known colonization routes (South genetic cluster: Hanover (Germany), Skåne (Sweden), Uppsala), North genetic cluster: Umeå (Sweden), Luleå (Sweden)). Contrasts were computed on estimated marginal means. Significant p-values adjusted for false discovery rate in bold.

**Growth rate including boldness (time to leave shelter shelter) as predictor**

| *contrast* | *Variable* | *estimate* | *SE* | *df* | *t.ratio* | *p.value* |
|---|---|---|---|---|---|---|
| North - South | Tadpole | 0.745* | 0.280 | 620 | 2.658 | **0.008** |
| | Froglet | -0.070 | 0.073 | 618 | -0.964 | 0.336 |
| Tadpole - Froglet | North | 0.512* | 0.235 | 621 | 2.180 | **0.030** |
| | South | -0.304 | 0.170 | 617 | -1.792 | 0.074 |

**Growth rate including exploration (mean distance to shelter) as predictor**

| *contrast* | *Variable* | *estimate* | *SE* | *df* | *t.ratio* | *p.value* |
|---|---|---|---|---|---|---|
| North - South | Tadpole | -0.117 | 0.075 | 476 | -1.565 | **0**.12 |
| | Froglet | 0.094 | 0.100 | 474 | 0.931 | 0.35 |
| Tadpole - Froglet | North | -0.263* | 0.108 | 475 | -2.438 | **0.015** |
| | South | -0.052 | 0.064 | 474 | -0.811 | 0.42 |

**Table S3. Trends of growth rate and behavior association by region and developmental stage.** Pairwise contrasts of slopes for the relationship between behaviors and developmental growth across regions and develomental stages in moor frog tadpoles and froglets originating from five different regions across a 1700 latitudinal gradient and raised in a common garden experiment (Hanover (Germany), Skåne (Sweden), Uppsala), Umeå (Sweden), Luleå (Sweden)). Contrasts were computed on estimated marginal means. Significant p-values adjusted for false discovery rate in bold.

**Boldness (time to leave shelter shelter)**

| *Dev. stage* | *region* | *Trend* | *SE* | *df* | *t.ratio* | *p.value* |
|---|---|---|---|---|---|---|
| Tadpole | Hanover | -0.376 | 0.268 | 603.004 | -1.406 | 0.229 |
| | Skåne | 0.886 | 0.351 | 605.509 | 2.526 | 0.059 |
| | Uppsala | 0.790* | 0.263 | 603.603 | 3.009 | **0.027** |
| | Umeå | -0.282 | 0.240 | 607.843 | -1.176 | 0.267 |
| | Luleå | -1.089 | 0.743 | 607.078 | -1.466 | 0.229 |
| Froglet | Germany | 0.142 | 0.111 | 603.004 | 1.275 | 0.254 |
| | Skåne | 0.115 | 0.066 | 606.007 | 1.755 | 0.199 |
| | Uppsala | -0.001 | 0.068 | 603.582 | -0.008 | 0.994 |
| | Umeå | 0.122 | 0.078 | 607.639 | 1.562 | 0.229 |
| | Luleå | 0.163 | 0.092 | 604.468 | 1.772 | 0.199 |

**Activity (mean speed)**

| *Dev. Stage* | *region* | *Trend* | *SE* | *df* | *t.ratio* | *p.value* |
|---|---|---|---|---|---|---|
| Tadpole | Hanover | -0.031 | 0.257 | 461.015 | -0.121 | 0.944 |
| | Skåne | 0.037 | 0.141 | 465.095 | 0.260 | 0.944 |
| | Uppsala | 0.005 | 0.071 | 468.702 | 0.070 | 0.944 |
| | Umeå | -0.023 | 0.054 | 468.252 | -0.433 | 0.944 |
| | Luleå | 0.007 | 0.052 | 461.557 | 0.140 | 0.944 |
| Froglet | Germany | -1.793 | 0.931 | 461.015 | -1.927 | 0.182 |
| | Skåne | -0.126 | 0.317 | 462.168 | -0.397 | 0.944 |
| | Uppsala | 0.727* | 0.246 | 463.509 | 2.959 | **0.032** |
| | Umeå | -0.320 | 0.567 | 461.158 | -0.565 | 0.944 |

| | Luleå | 2.468 | 0.959 | 464.822 | 2.572 | 0.052 |
|---|---|---|---|---|---|---|

**Exploration (mean distance to shelter)**

| *Dev. Stage* | *region* | *Trend* | *SE* | *df* | *t.ratio* | *p.value* |
|---|---|---|---|---|---|---|
| Tadpole | Hanover | -0.191 | 0.124 | 461.016 | -1.541 | 0.493 |
| | Skåne | 0.035 | 0.064 | 463.134 | 0.549 | 0.834 |
| | Uppsala | 0.003 | 0.066 | 461.641 | 0.038 | 0.970 |
| | Umeå | -0.112 | 0.096 | 466.914 | -1.168 | 0.585 |
| | Luleå | -0.147 | 0.081 | 464.300 | -1.808 | 0.493 |
| Froglet | Germany | -0.047 | 0.163 | 461.016 | -0.286 | 0.861 |
| | Skåne | 0.031 | 0.083 | 461.396 | 0.375 | 0.861 |
| | Uppsala | 0.058 | 0.066 | 461.168 | 0.872 | 0.639 |
| | Umeå | 0.122 | 0.116 | 461.836 | 1.054 | 0.585 |
| | Luleå | 0.215 | 0.149 | 463.506 | 1.449 | 0.493 |

**Table S4. Independent contrasts of growth rate models by developmental stage.** Statistical contrasts of significant interaction effects between regions in models evaluating growth rate in moor frog tadpoles and froglets originating from five different regions across a 1700 latitudinal gradient and raised in a common garden experiment (Hanover (Germany), Skåne (Sweden), Uppsala), Umeå (Sweden), Luleå (Sweden)). Contrasts were computed on estimated marginal means. Significant p-values adjusted for false discovery rate in bold.

**Growth rate including boldness (time to leave shelter shelter) as predictor**

| *Dev.stage* | *contrast* | *estimate* | *SE* | *df* | *t.ratio* | *p.value* |
|---|---|---|---|---|---|---|
| Tadpole | Hanover - Skåne | 1.262 | 0.441 | 604.639 | 2.861 | **0.015** |
| | Hanover - Uppsala | 1.167 | 0.375 | 603.301 | 3.111 | **0.013** |
| | Hanover - Umeå | 0.094 | 0.359 | 605.423 | 0.262 | 0.828 |
| | Hanover - Luleå | -0.713 | 0.790 | 606.681 | -0.903 | 0.459 |
| | Skåne - Uppsala | -0.095 | 0.438 | 604.855 | -0.217 | 0.828 |
| | Skåne - Umeå | -1.168 | 0.425 | 606.321 | -2.749 | **0.015** |
| | Skåne - Luleå | -1.975 | 0.821 | 606.811 | -2.404 | **0.029** |
| | Uppsala - Umeå | -1.072 | 0.356 | 605.743 | -3.015 | **0.013** |
| | Uppsala - Luleå | -1.879 | 0.788 | 606.744 | -2.385 | **0.029** |
| | Umeå - Luleå | -0.807 | 0.781 | 607.154 | -1.034 | 0.431 |
| Froglet | Hanover - Skåne | -0.027 | 0.129 | 603.843 | -0.206 | 0.950 |
| | Hanover - Uppsala | -0.142 | 0.131 | 603.164 | -1.090 | 0.690 |
| | Hanover - Umeå | -0.020 | 0.136 | 604.736 | -0.149 | 0.950 |
| | Hanover - Luleå | 0.021 | 0.144 | 603.614 | 0.145 | 0.950 |
| | Skåne - Uppsala | -0.116 | 0.095 | 604.802 | -1.221 | 0.690 |
| | Skåne - Umeå | 0.006 | 0.102 | 606.998 | 0.062 | 0.950 |
| | Skåne - Luleå | 0.048 | 0.113 | 605.010 | 0.421 | 0.950 |
| | Uppsala - Umeå | 0.122 | 0.104 | 606.059 | 1.178 | 0.690 |
| | Uppsala - Luleå | 0.163 | 0.114 | 604.158 | 1.426 | 0.690 |
| | Umeå - Luleå | 0.041 | 0.120 | 605.916 | 0.342 | 0.950 |

**Growth rate including activity (mean speed) as predictor**

| *Dev. stage* | *contrast* | *estimate* | *SE* | *df* | *t.ratio* | *p.value* |
|---|---|---|---|---|---|---|
| Tadpole | Hanover - Skåne | -0.068 | 0.293 | 462.077 | -0.231 | 0.979 |
| | Hanover - Uppsala | -0.036 | 0.267 | 461.892 | -0.136 | 0.979 |

| | | | | | | |
|---|---|---|---|---|---|---|
| | Hanover - Umeå | -0.008 | 0.263 | 461.840 | -0.030 | 0.979 |
| | Hanover - Luleå | -0.038 | 0.262 | 461.037 | -0.147 | 0.979 |
| | Skåne - Uppsala | 0.032 | 0.158 | 466.156 | 0.200 | 0.979 |
| | Skåne - Umeå | 0.060 | 0.151 | 466.373 | 0.398 | 0.979 |
| | Skåne - Luleå | 0.029 | 0.150 | 464.726 | 0.195 | 0.979 |
| | Uppsala - Umeå | 0.028 | 0.089 | 468.999 | 0.317 | 0.979 |
| | Uppsala - Luleå | -0.002 | 0.088 | 467.273 | -0.026 | 0.979 |
| | Umeå - Luleå | -0.031 | 0.075 | 468.194 | -0.409 | 0.979 |
| Froglet | Hanover - Skåne | -1.667 | 0.983 | 461.139 | -1.696 | 0.113 |
| | Hanover - Uppsala | -2.520 | 0.963 | 461.191 | -2.618 | **0.032** |
| | Hanover - Umeå | -1.473 | 1.090 | 461.054 | -1.351 | 0.197 |
| | Hanover - Luleå | -4.261 | 1.337 | 463.121 | -3.188 | **0.015** |
| | Skåne - Uppsala | -0.853 | 0.401 | 462.688 | -2.127 | 0.068 |
| | Skåne - Umeå | 0.194 | 0.649 | 461.404 | 0.300 | 0.765 |
| | Skåne - Luleå | -2.594 | 1.010 | 464.588 | -2.567 | **0.032** |
| | Uppsala - Umeå | 1.047 | 0.618 | 461.556 | 1.695 | 0.113 |
| | Uppsala - Luleå | -1.741 | 0.990 | 464.746 | -1.758 | 0.113 |
| | Umeå - Luleå | -2.788 | 1.114 | 463.978 | -2.502 | **0.032** |

**Table S5. Independent contrasts of growth rate models by region.** Statistical contrasts of significant interaction effects between developmental stages in models evaluating growth rate in moor frog tadpoles and froglets originating from five different regions across a 1700 latitudinal gradient and raised in a common garden experiment (Hanover (Germany), Skåne (Sweden), Uppsala), Umeå (Sweden), Luleå (Sweden)). Contrasts were computed on estimated marginal means. Significant p-values adjusted for false discovery rate in bold.

**Growth rate including boldness (time to leave shelter shelter) as predictor**

| *contrast* | *region* | *estimate* | *SE* | *df* | *t.ratio* | *p.value* |
|---|---|---|---|---|---|---|
| Tadpole - Froglet | Hanover | -0.518 | 0.290 | 603.004 | -1.788 | 0.074 |
| | Skåne | 0.770 | 0.356 | 604.822 | 2.163 | **0.031** |
| | Uppsala | 0.791 | 0.272 | 603.876 | 2.912 | **0.004** |
| | Umeå | -0.404 | 0.251 | 605.675 | -1.609 | 0.108 |
| | Luleå | -1.252 | 0.748 | 606.909 | -1.673 | 0.095 |

**Growth rate including activity (mean speed) as predictor**

| *contrast* | *region* | *estimate* | *SE* | *df* | *t.ratio* | *p.value* |
|---|---|---|---|---|---|---|
| Tadpole - Froglet | Hanover | 1.762 | 0.966 | 461.015 | 1.825 | 0.069 |
| | Skåne | 0.163 | 0.348 | 464.109 | 0.467 | 0.641 |
| | Uppsala | -0.722 | 0.256 | 464.031 | -2.822 | **0.005** |
| | Umeå | 0.297 | 0.569 | 461.078 | 0.522 | 0.602 |
| | Luleå | -2.460 | 0.961 | 464.766 | -2.561 | **0.011** |

**Growth rate including exploration (mean distance to shelter) as predictor**

| *contrast* | *region* | *estimate* | *SE* | *df* | *t.ratio* | *p.value* |
|---|---|---|---|---|---|---|
| Tadpole - Froglet | Hanover | -0.144 | 0.205 | 461.016 | -0.704 | 0.482 |
| | Skåne | 0.004 | 0.104 | 461.621 | 0.038 | 0.970 |
| | Uppsala | -0.055 | 0.094 | 461.393 | -0.590 | 0.555 |
| | Umeå | -0.234 | 0.151 | 465.553 | -1.552 | 0.121 |
| | Luleå | -0.362 | 0.170 | 465.554 | -2.129 | **0.034** |

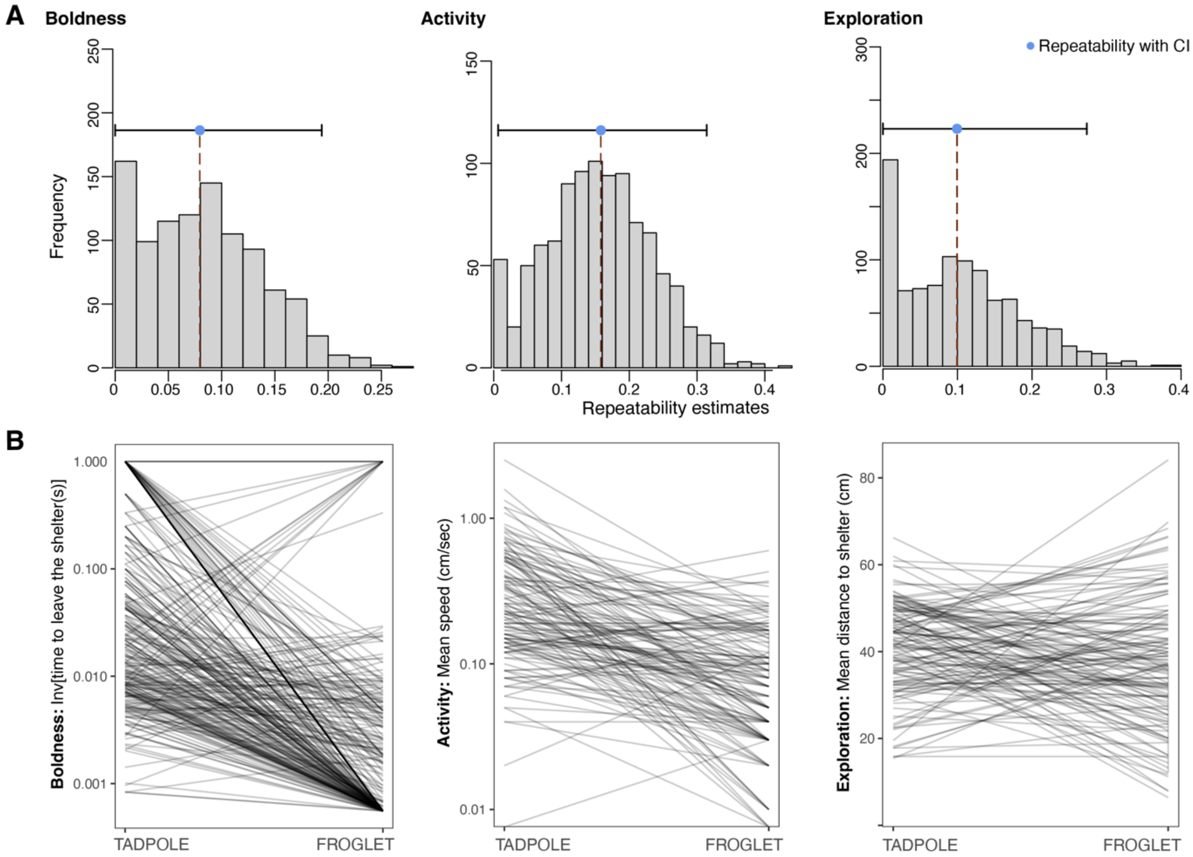


**Figure S1. Consistency of individual behaviour across developmental stages.** (A) Distribution of adjusted repeatability estimates of measurements in time to emerge from the shelter, activity and explorations taken as tadpole and froglet to each individual. Blue dot indicates mean adjusted repeatability with confidence intervals for each behavioural trait. (B) Individual differences in measurements of time to emerge from the shelter, activity and exploration taken to individuals as tadpole and froglet developmental stage. Lines represents measurements for each individual.

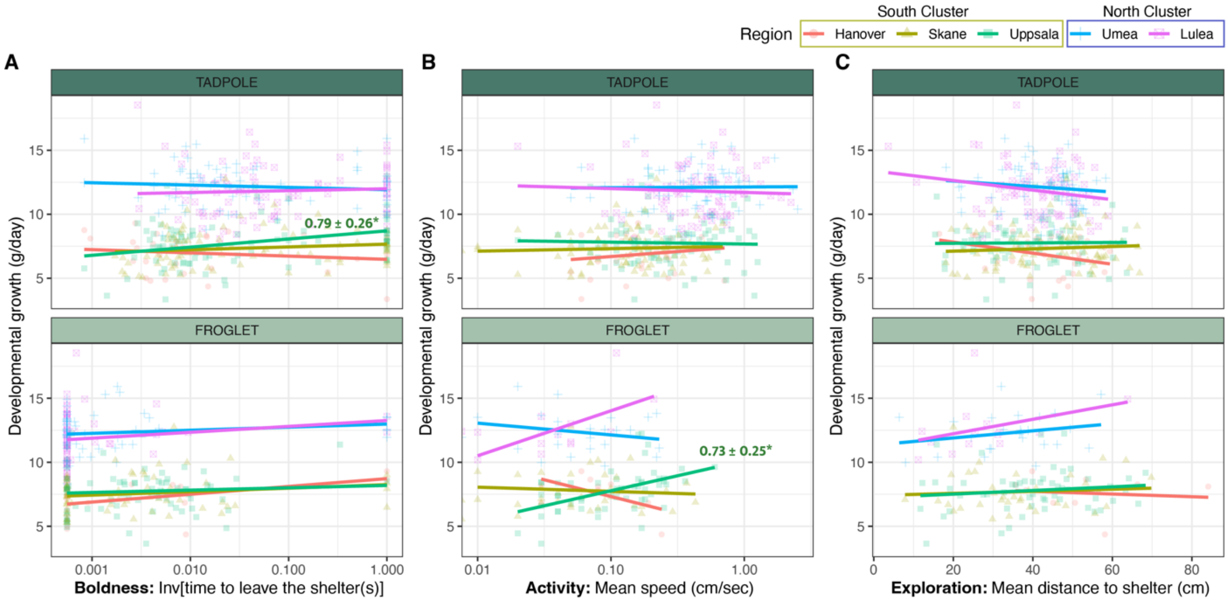


**Figure S2. Association between behaviour and life history in moor frogs by region.** Relationships between (A) boldness, (B) activity and (C) exploration and developmental growth (mass/time at Gosner stage 42) in moor frog tadpoles (top row) and froglets (bottom row) sampled in several regions across a 1700 latitudinal gradient and raised in a common garden experiment (South genetic cluster: Hanover (Germany), red dot; Skåne (Sweden), yellow triangle; Uppsala, green square); North genetic cluster: Umeå (Sweden), blue cross; Luleå (Sweden), pink crossed square). Trend lines are displayed for the relationship between each behaviour and developmental growth for values at the all regions. Text indicates estimated marginal means and standard error for significant trends in post-hoc contrasts of statistical models (see Table S5 for extended information on trends for each region).
